# Distinct roles of LARP1 and 4EBP1/2 in regulating translation and stability of 5′TOP mRNAs

**DOI:** 10.1101/2023.05.22.541712

**Authors:** Tobias Hochstoeger, Panagiotis Papasaikas, Ewa Piskadlo, Jeffrey A. Chao

## Abstract

A central mechanism of mTOR complex 1 (mTORC1) signaling is the coordinated translation of ribosomal protein and translation factor mRNAs mediated by the 5′-terminal oligopyrimidine motif (5′TOP). Recently, La-related protein 1 (LARP1) has been proposed to be the specific regulator of 5′TOP mRNA translation downstream of mTORC1, while eIF4E-binding proteins (4EBP1/2) were suggested to have a general role in repression. Here, we employ single-molecule translation site imaging of 5′TOP and canonical mRNAs to study the translational dynamics of single mRNAs in living cells. Our data reveals that 4EBP1/2 has a dominant role in translation repression of both 5′TOP and canonical mRNAs during pharmacological inhibition of mTOR. In contrast, we find that LARP1 selectively protects 5′TOP mRNAs from degradation in a transcriptome-wide analysis of mRNA half-lives. Our results clarify the roles of 4EBP1/2 and LARP1 in regulating 5′TOP mRNAs and provides a framework to further study how these factors control cell growth during development and disease.

## Introduction

For cellular homeostasis, ribosome biogenesis needs to be tightly coupled to nutrient availability. In eukaryotic cells, mTORC1 is the central signaling hub that integrates nutrient cues to match cell growth by stimulating or inhibiting ribosome biogenesis (1, 2). When nutrients are available, active mTORC1 promotes translation by the phosphorylation of key substrates, S6 kinases and 4EBP1/2, stimulating eIF4F assembly and translation. When nutrients are limited, mTORC1 substrates are dephosphorylated, allowing 4EBP1/2 to bind and sequester the cap-binding protein eIF4E, thereby inhibiting mRNA translation. In addition, the La-related protein 1 (LARP1) has been described as a direct mTORC1 substrate and translational regulator (3–5). While mTORC1-dependent translation regulation acts on all mRNAs via multiple routes, it exerts a much more rapid and pronounced effect on ribosomal protein and translation factor mRNAs (∼100 mRNAs) that carry a 5′-terminal oligopyrimidine motif (5′TOP, 4-15 pyrimidines) directly adjacent to the 5′ cap (6).

While it has been well established that the 5′TOP motif is both essential and sufficient for rapid mTORC1-mediated translational regulation (7, 8), the underlying molecular mechanism has been challenging to resolve (9). Both 4EBP1/2 and LARP1 have been found to mediate 5′TOP translational repression, as loss of either factor partially relieved 5′TOP translational repression in cells acutely treated with the mTOR inhibitor Torin1 (4, 5, 10, 11). Importantly, a crystal structure of LARP1 bound to both the 5′ cap and first five nucleotides of a 5′TOP oligo provided mechanistic insight into how LARP1 could specifically repress 5′TOP mRNAs upon mTORC1 inhibition, leading to a model in which dephosphorylated LARP1 binds the 5′ end of 5′TOP mRNAs to prevent assembly of the eIF4F complex (12). Mechanistic insight for 4EBP1/2 mediated translational regulation is still lacking, but could be mediated by interactions of eIF4F with the cap-adjacent nucleotides of the 5′TOP motif (13, 14).

An additional layer of ribosome biogenesis control is the pool of 5′TOP mRNAs available for translation, which are among the most highly expressed and stable transcripts in eukaryotic cells (15, 16). LARP1 has been found to associate with PABPC1 and inhibit deadenylation of mRNA transcripts (17–20). It is currently unclear how LARP1 is recruited to the mRNAs it stabilizes, the importance of the 5′TOP motif for target selection, and the relevance of mTORC1 activity in this process. Crosslinking studies have found LARP1 bound to thousands of mRNAs, a subset of which are bound more upon mTORC1 inhibition (including, but not limited to 5′TOP mRNAs) (21–23). In contrast, polyA-tail sequencing in mTORC1 active cells have found LARP1 to inhibit mRNA deadenylation globally, with stronger protection of 5′TOP transcripts (24).

In this study, we sought to clarify the roles of LARP1 and 4EBP1/2 in regulating the translation and stability of 5′TOP mRNAs. Direct measurements of translation of 5′TOP and non-5′TOP (canonical) mRNAs using single-molecule SunTag imaging revealed a dominant role of 4EBP1/2 in in mediating 5′TOP translational repression. In contrast, we find a highly selective role of LARP1 in protecting 5′TOP mRNAs from degradation by measuring genome-wide changes in mRNA half-lives using metabolic mRNA labeling (SLAMseq). Our study provides insights into the distinct roles of LARP1 and 4EBP1/2 in mediating 5′TOP translational regulation, and a framework for further investigations into the mechanisms by which these factors regulate cell growth under normal physiological conditions and disease.

## Results

### Single-molecule imaging of translation during mTOR inhibition

To study the regulation of translation during mTOR inhibition, we engineered a HeLa cell line that expresses fluorescent proteins for single-molecule imaging of mRNA (NLS-MCP-Halo) and translation (scFV-GFP), together with the reverse tetracycline-controlled transactivator to enable induction of reporter mRNAs (25). Into this cell line, we integrated two different constructs into a single genomic locus under the control of a doxycycline-inducible promoter. The reporter mRNAs were identical except for their 5′UTR, where one contains a 5′TOP motif as part of the full-length RPL32 5′UTR, and the other has a non-5′TOP canonical 5′UTR. The coding sequence encodes 24 GCN4 epitope tags for translation site imaging (SunTag, (26)) followed by Renilla luciferase for bulk measurements of translation and the FKBP12-derived destabilization domain to reduce the accumulation of mature proteins (27). In addition, the 3′UTR contains 24 MS2 stem-loops for mRNA imaging (Fig. 1A). The 5′-end of both reporter mRNAs was sequenced in order to determine the transcription start sites. All 5′TOP transcripts contained a 5′TOP motif and the canonical transcripts initiated with AGA, which is similar to the most common transcription start site (Fig. S1) (28). To inhibit mTOR, we used the ATP-competitive inhibitor Torin1, which has been widely used to study the translation of 5′TOP mRNAs. For both the 5′TOP and canonical mRNA reporter cell lines, Torin1 treatment for 1hr resulted in inhibition of mTORC1 as seen by 4EBP1 dephosphorylation, which is also consistent with previous results (Fig. S2) (21).

**Fig. 1.**
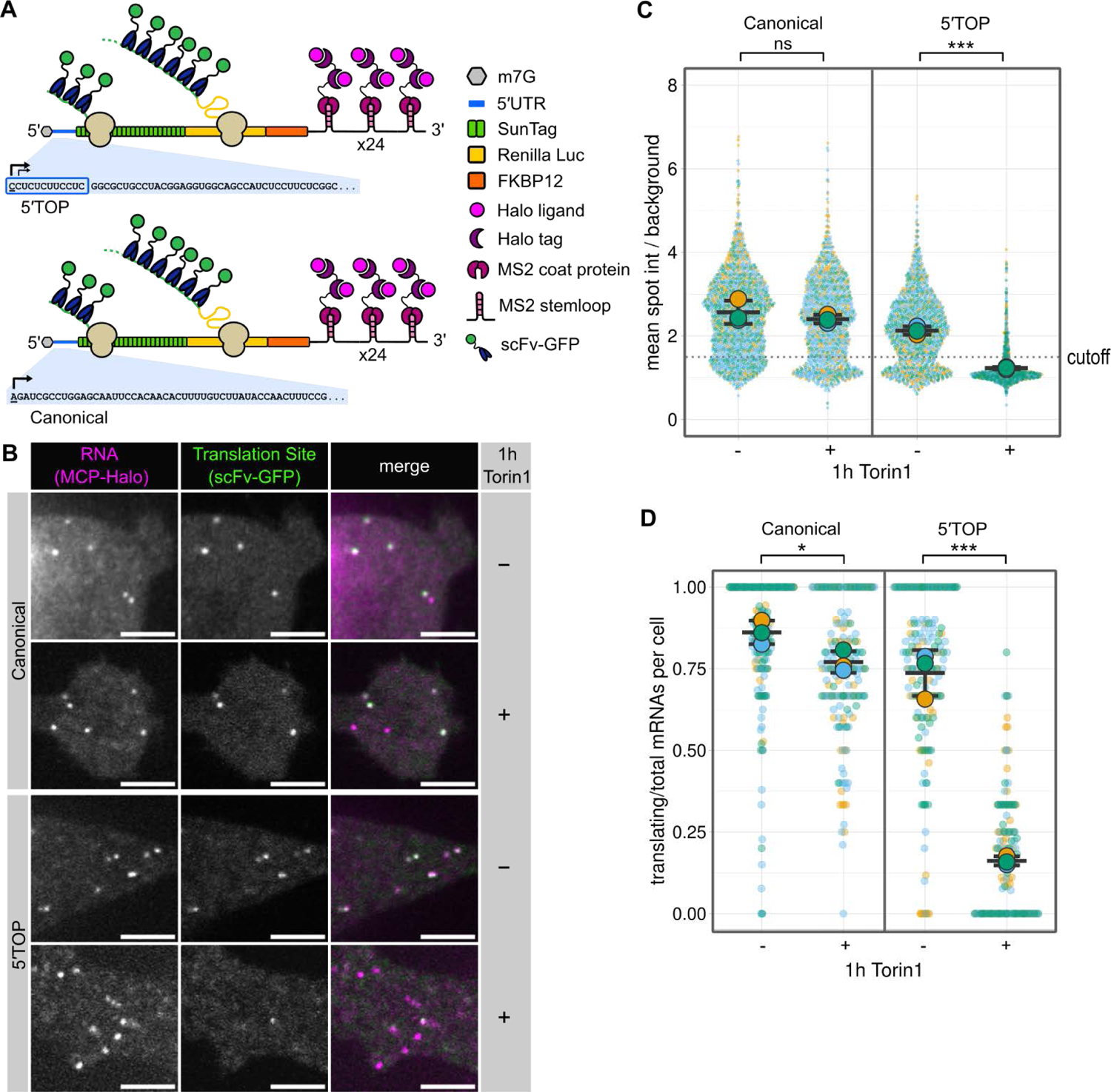
Single-molecule imaging recapitulates 5′TOP translational repression. (**A**) Schematic representation of reporter mRNAs for single molecule imaging of translation. The 5′TOP reporter contains the full-length RPL32 5′UTR, whereas the canonical reporter has a control 5′UTR of similar length. Black arrows indicate transcription start sites. (**B**) Representative images of canonical and 5′TOP reporter mRNAs (MCP-Halo foci, magenta) undergoing translation (scFv-GFP foci, green) in absence or presence of mTOR inhibitor Torin 1 (250nM, 1h). Scale bars = 5 µm. (**C**) Translation site intensities of canonical and 5′TOP reporter mRNAs quantified in absence and presence of Torin 1 (250nM, 1h). SunTag intensities are plotted for all mRNAs (colored circles) overlaid with the mean ± SD (≥1089 mRNAs per condition, n=3). (**D**) Fraction of mRNAs undergoing translation quantified per cell for canonical and 5′TOP reporter in absence or presence of Torin 1 (250nM, 1h). Values are plotted for each cell (colored circles) overlaid with the mean ± SD (≥162 cells per condition, n=3). For statistics, unpaired t tests were performed, with statistical significance claimed when p < 0.05 (ns = not significant, * = p < 0.05, *** = p < 0.001).

To observe the effect of mTORC1 inhibition on translation, we induced expression of the reporter mRNAs in both cell lines and imaged them in either the presence or absence of Torin1. Treatment with Torin1 was found to strongly repress translation of most 5′TOP transcripts as seen by the disappearance of scFv-GFP spots that co-localized with mRNA spots, whereas the canonical mRNAs were largely unaffected (Fig. 1B). For quantification of single mRNAs and their translation, we employed a high-throughput analysis pipeline that tracks individual mRNAs and measures the corresponding SunTag intensities. mRNA trajectories were determined using single-particle tracking of MCP-Halo spots, and scFv-GFP intensities at those same coordinates were quantified for each mRNA as background corrected mean spot intensity (SunTag intensity). Using this analysis pipeline, we quantified the translation of >1000 mRNAs for both the 5′TOP and canonical mRNA cell lines (Fig. 1C), which revealed a broad distribution of SunTag intensities for both types of transcripts indicating a heterogeneity of ribosomes engaged in translation of individual transcripts (26, 29). The average SunTag intensity for the 5′TOP mRNAs was slightly lower compared to the canonical mRNAs indicating fewer ribosomes engaged in translation when mTOR is active (Fig. 1C). The mean SunTag intensity for the 5′TOP mRNAs decreased markedly upon Torin1 treatment, whereas the mean SunTag intensity of the canonical mRNAs decreased only slightly.

While changes in SunTag intensity indicate differences in ribosome number, translation site imaging can also be used to quantify the fraction of transcripts actively translating within a cell. Puromycin treatment, which inhibits translation due to premature termination, was used to measure SunTag spot intensities in the absence of translation in order to calibrate a threshold for identifying translating mRNAs (>1.5-fold over background, Fig. S3). Quantifying translation as the fraction of translating mRNAs per cell revealed slightly fewer translating 5′TOP mRNAs (mean: 74%) compared to the canonical mRNAs (mean: 86%) when mTOR is active (Fig. 1D). Upon 1hr Torin1 treatment, the fraction of translating 5′TOP mRNAs per cell decreased drastically (mean: 16%), though many cells retained a minor fraction of translating 5′TOP mRNAs. In contrast, the fraction of translating canonical mRNAs decreased only slightly upon Torin1 treatment (mean: 77%). To determine whether the remaining fraction of translating 5′TOP mRNAs after 1hr Torin1 treatment represented stalled ribosomes, Torin1 treated cells were co-treated with harringtonine, which stalls ribosome at the start codon to deplete elongating ribosomes. Addition of harringtonine abolished the remaining translation sites in the Torin1-treated 5′TOP cell line within 10 min (Fig. S4), demonstrating that the low number of 5′TOP mRNAs that co-localize with SunTag signal are still actively translating. Taken together, our data captures both inter- and intracellular variability in the translation of canonical and 5′TOP mRNAs in the presence and absence of Torin1, providing direct translation measurements uncoupled from transcriptional regulation or mRNA stability.

### LARP1 knockout partially rescues translation of 5′TOP mRNAs during Torin1 treatment

Recently, LARP1 has been found to specifically bind the 5′TOP motif in an mTOR-dependent manner to regulate translation (3, 12, 21). To further investigate the role of LARP1 in translational inhibition of 5′TOP mRNAs during mTOR inhibition, we generated LARP1 CRISPR-Cas9 knockouts in the 5′TOP and canonical mRNA cell lines. Genomic DNA sequencing confirmed frameshift mutations in all alleles of LARP1 exon 4 that are upstream of any domain of known function (aa205-aa240) (Fig. S5A and B). Loss of LARP1 protein in the knockout cell lines was confirmed by western blot analysis using two LARP1 antibodies targeting either the N- or C-terminal regions, which did not detect alternative LARP1 isoforms (Fig. S5C). Importantly, loss of LARP1 did not disrupt the regulation of other mTORC1 targets, as seen by inhibition of 4EBP1 phosphorylation upon 1hr Torin1 treatment (Fig. S5D). Consistent with earlier reports in HEK cells, deletion of LARP1 in HeLa cells resulted in decreased cell proliferation (4, 5).

Following the validation of the LARP1 KO cell lines, we quantified the translation of 5′TOP and canonical mRNAs (>600 mRNAs per condition) in the absence of LARP1 (Fig. S6A). Analysis of SunTag intensities of the canonical mRNAs revealed similar translation levels in the LARP1 KO compared to WT. Upon 1hr Torin1 treatment, canonical mRNAs decreased slightly in mean SunTag intensity, whereas the 5′TOP mRNAs decreased more strongly (Fig. 2A). Calculating the fraction of translating mRNAs per cells revealed that the canonical mRNAs show a mild response to Torin1 in the absence of LARP1 (mean untreated: 87%, mean 1hr Torin1: 75%, Fig. 2B), mirroring the response observed for the canonical mRNAs in LARP1 WT cells. Interestingly, the 5′TOP mRNAs in LARP1 KO cells displayed a partial rescue of translational inhibition upon Torin1 treatment (mean untreated: 79%, mean 1hr Torin1: 41%) compared to LARP1 WT cells (mean 1hr Torin1: 16%). The incomplete rescue of 5′TOP mRNA translation in the absence of LARP1 suggests the existence of additional *trans*-regulatory factors in mediating 5′TOP translational repression.

**Fig. 2.**
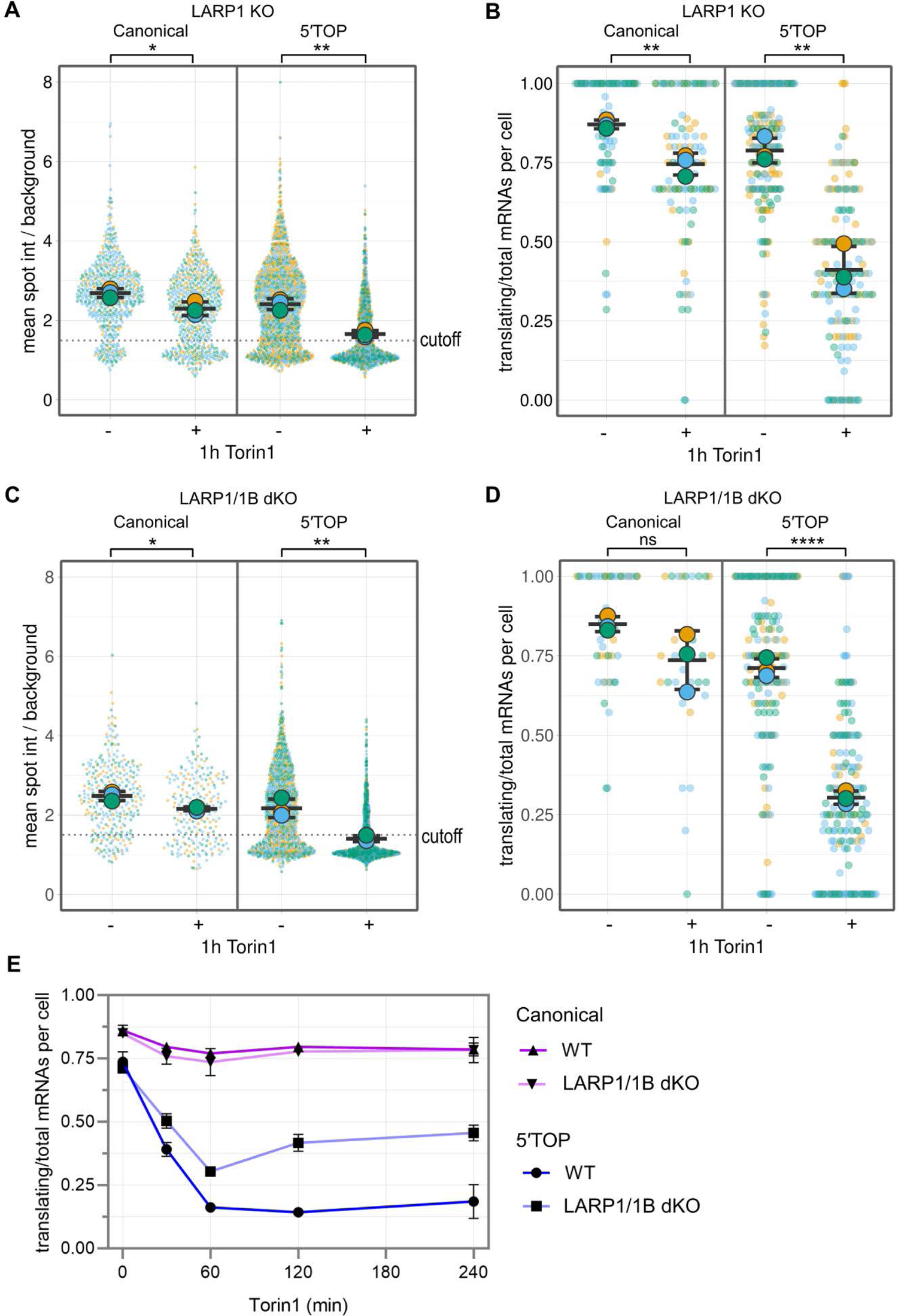
Loss of LARP1 partially alleviates 5′TOP translational repression during Torin1 treatment. (**A**) Quantification of translation site intensities in LARP1 KO cells ± Torin1 (250nM, 1h). SunTag intensities are plotted for all mRNAs overlaid with the mean ± SD (≥652 mRNAs per condition, n=3). (**B**) Fraction of mRNAs undergoing translation quantified per cell in LARP1 KO cells ± Torin1 (250nM, 1h). Values are plotted for each cell (colored circles) overlaid with the mean ± SD (≥91 cells per condition, n=3). (**C**) Quantification of translation site intensities in LARP1/1B double KO cells ± Torin1 (250nM, 1h). SunTag intensities are plotted for all mRNAs (colored circles) overlaid with the mean ± SD (≥218 mRNAs per condition, n=3). (**D**) Fraction of mRNAs undergoing translation quantified per cell in LARP1/1B double KO cells ± Torin1 (250nM, 1h). Values are plotted for each cell (colored circles) overlaid with the mean ± SD (≥30 cells per condition, n=3). For statistics, unpaired t tests were performed, with statistical significance claimed when p < 0.05 (ns = not significant, * = p < 0.05, ** = p < 0.01, **** = p < 0.0001. (**E**) Time course of fraction of translating mRNAs per cell for canonical and 5′TOP reporter cell lines in the presence (WT) or absence of LARP1/1B (dKO). Cells were treated with 0, 30, 60, 120, and 240 min Torin1. Values are plotted as the mean ± SEM (≥30 cells per condition, n=3).

One possible trans-acting factor that could repress 5′TOP mRNAs in the absence of LARP1 is the homolog LARP1B (also called LARP2), which shares the DM15 domain that binds the 5′TOP motif, though it is lowly expressed in HeLa cells. To test this possibility, we used CRISPR-Cas9 to generate knockouts of LARP1B in the LARP1 knockout background (Fig. S7A). Genomic DNA sequencing of the edited alleles identified frameshift mutations in all alleles of LARP1B exon 4, and PCR confirmed loss of wildtype LARP1B mRNA (Fig. S7B and C). Western blot analysis confirmed unperturbed mTORC1 signaling in the LARP1/1B double KO (dKO) cells, as seen by 4EBP1 dephosphorylation upon 1hr Torin1 treatment (Fig. S7D).

Following the validation of the LARP1/1B dKO cell lines, we quantified the translation of 5′TOP and canonical mRNAs (>200 mRNAs per condition) in the absence of LARP1/1B (Fig. S6B). Both the distribution of SunTag intensities (Fig 2C) and the fraction of translating canonical and 5′TOP mRNAs per cell (Fig. 2D) responded similarly to Torin1 treatment (1hr) as obtained for LARP1 KO cells, suggesting that LARP1B does not affect 5′TOP translational regulation in our HeLa cells. These results are consistent with an earlier study in HEK cells that also found that translational repression of endogenous 5′TOP transcripts could not be rescued further by combinatory deletion of LARP1/1B (30), arguing against functional redundancy of LARP1 and LARP1B.

While we did not observe a rescue of 5′TOP translation when cells were treated with Torin1 for 1hr, we could not exclude the possibility that the effect of loss of LARP1/1B might be more pronounced at other time points. To characterize the kinetics of Torin1 translational inhibition, we performed SunTag imaging of the canonical and 5′TOP cell lines at additional time points 30 min, 2hr, and 4hr (Fig. 2E). For both WT and LARP1/1B dKO cells, the canonical mRNAs showed a gradual decrease in translation during the first hour of Torin1 treatment that remained constant at later time points. To test whether prolonged mTOR inhibition is required to repress canonical mRNA translation, we quantified translation of canonical mRNAs in WT cells treated with Torin1 for 24hr. Interestingly, the majority of canonical mRNAs remained translating at this longer timepoint (Fig. S8). 5′TOP mRNAs in WT cells decreased in translation within 1hr of Torin1 to a minor fraction of translating mRNAs (30min Torin1: 39%, 1hr Torin1: 16%) and remained at this level at the 2hr (14%) and 4hr (19%) timepoints (Fig. 2E). In LARP1/1B dKO cells, translation of 5′TOP mRNAs also decreased with no change in the timing of repression, but a decrease in its extent (30min Torin1: 50%, 1hr Torin1: 30%), however at the 2hr (42%) and 4hr (46%) timepoints, we observed a slight increase in translation. These results suggest that while LARP1 may contribute to translation repression of 5ʹTOP mRNAs, it is not the dominant regulatory factor during mTOR inhibition.

### 4EBP1/2 knockout rescues translation of 5′TOP mRNAs during Torin1 treatment

The eIF4E-binding proteins 1 and 2 (4EBP1/2) are thought to generally repress translation during mTORC1 inhibition but have also been previously implicated in specifically affecting 5′TOP mRNAs (10). Using lentiviral infection, stable shRNA-mediated knockdown (KD) cell lines were generated in the wildtype and LARP1/1B dKD background for both 5′TOP and canonical mRNA cell lines. Western blot analysis confirmed the depletion of 4EBP1/2 levels in all four cell lines (Fig. S9A and B). Furthermore, the dephosphorylation of RPS6 and residual 4EBP1 upon 1hr Torin1 indicated that mTORC1 signaling was unperturbed in the 4EBP1/2 dKD cell lines (Fig. S9C and D).

Having validated the 4EBP1/2 dKD cell lines, we measured the translation of canonical and 5′TOP mRNAs in the absence of 4EBP1/2 (Fig. S10). In untreated cells, the reduction of 4EBP1/2 resulted in increased SunTag intensities for both canonical and 5′TOP mRNAs in cell lines with WT LARP1 (Fig. 3A, >1000 mRNAs per condition) and LARP1/1B dKO (Fig. 3B, >700 mRNAs per condition) compared to cells with wildtype levels of 4EBP1/2 (Fig. 1C). This suggested that when mTOR is active, 4EBP1/2 can still weakly repress translation initiation presumably through fluctuations in mTOR signaling during cell growth. Surprisingly, when 4EBP1/2 were depleted, the SunTag intensities of both canonical and 5′TOP mRNAs were not reduced upon 1hr Torin1 treatment in cells with WT LARP1 (Fig. 3A) or LARP1/1B dKO (Fig. 3B). Analyzing the fraction of translating canonical mRNAs revealed no change in translation upon Torin1 treatment for cells with WT LARP1 (Fig. 3C) and LARP1/1B dKO (Fig. 3D), in contrast to the previously observed mild decrease in translation of canonical mRNAs upon 1hr Torin1 (Fig 1D). Importantly, when 4EBP1/2 levels are depleted, the fraction of translating mRNAs of 5′TOP mRNAs is also similar during Torin1 treatment of cells with WT LARP1 (Fig. 3B, untreated: 77%, treated: 70%) and LARP1/1B dKO (Fig. 3D, untreated: 79%, treated: 77%). To exclude the possibility that translational repression is delayed in the absence of 4EBP1/2, we investigated the kinetics of mTOR inhibition in the 4EBP1/2 dKD cell lines (Fig. 3E), which revealed that canonical and 5′TOP mRNAs remain similarly insensitive to Torin1 treatment at prolonged Torin1 treatment (2hr, 4hr). These experiments indicate that despite the difference in magnitude of translation repression during Torin1 treatment, 4EBP1/2 is responsible for the weak inhibition of canonical mRNAs and the stronger inhibition of 5′TOP mRNAs. Our data supports a model where 5′TOP mRNAs are intrinsically more sensitive to 4EBP1/2 mediated translational regulation, which results in a minor difference in translation when mTOR is active, and a pronounced difference in translation when mTOR is inhibited.

**Fig. 3.**
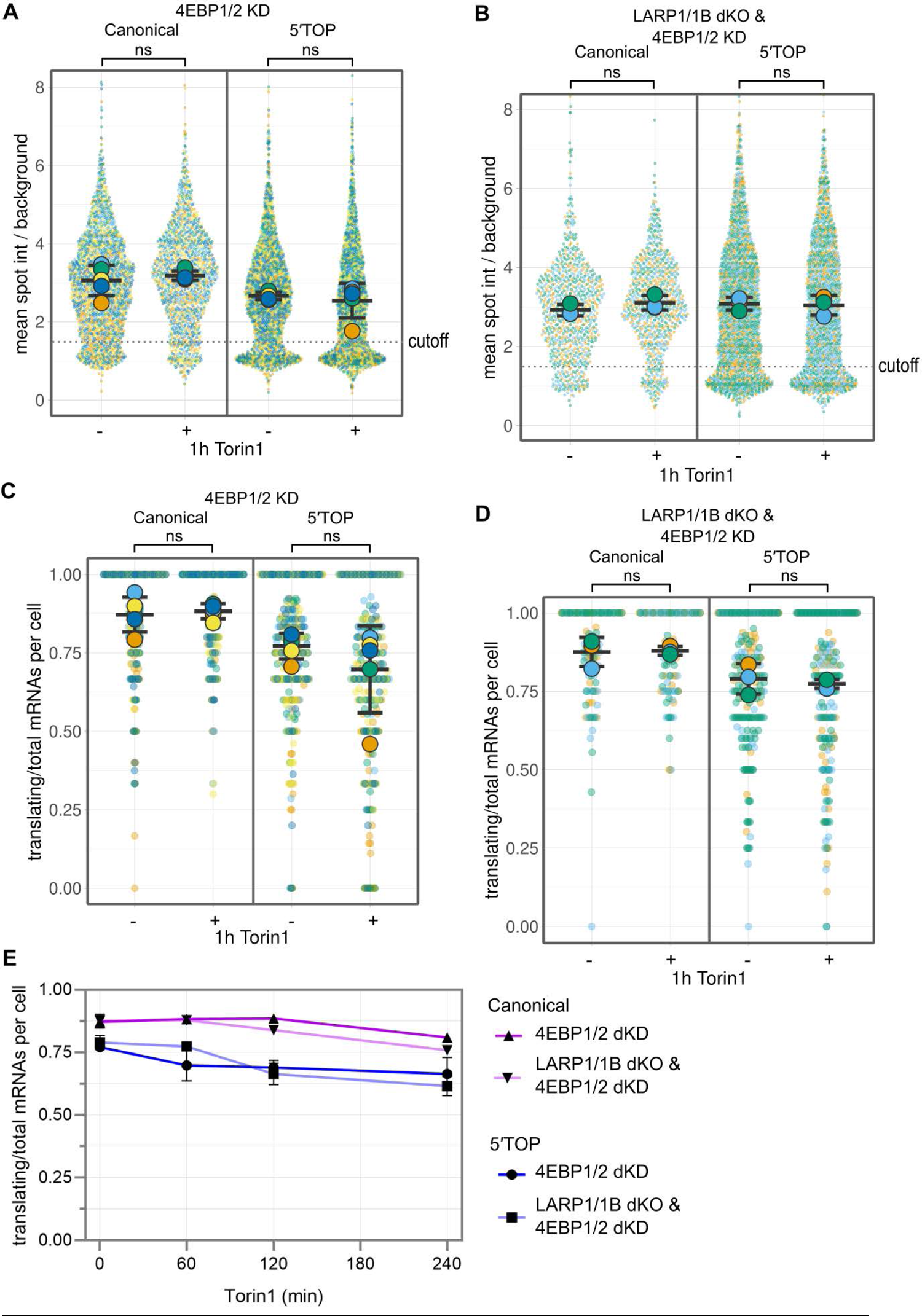
Loss of 4EBP1/2 is sufficient to alleviate 5′TOP translational repression during Torin1 treatment. (**A**) Quantification of translation site intensities of reporter mRNAs in 4EBP1/2 dKD cells ± Torin1 (250nM, 1h). SunTag intensites are plotted for all mRNAs (colored circles) overlaid with the mean ± SD (≥1344 mRNAs per condition, n=5). (**B**) Fraction of mRNAs undergoing translation quantified per cell for reporter mRNAs in 4EBP1/2 dKD cells ± Torin1 (250nM, 1h). Values are plotted for each cell (colored circles) overlaid with the mean ± SD (≥204 cells per condition, n=5). (**C**) Quantification of translation site intensities of reporter mRNAs in LARP1/1B dKO_4EBP1/2 dKD cells ± 1hr Torin1 (250nM). SunTag intensities are plotted for all mRNAs overlaid with the mean ± SD (≥783 mRNAs per condition, n=3). (**D**) Fraction of mRNAs undergoing translation quantified per cell of reporter mRNAs in LARP1/1B dKO_4EBP1/2 dKD cells ± Torin1 (250nM, 1h). Values are plotted for each cell (colored circles) overlaid with the mean ± SD (≥97 cells per condition, n=3). For statistics, unpaired t tests were performed, with statistical significance claimed when p < 0.05 (ns = not significant). (**E**) Time course of fraction translating mRNAs per cell for canonical and 5′TOP reporters in 4EBP1/2 dKD cells with WT LARP1 or in combination with LARP1/1B dKO. Cells were treated with 0, 30, 60, 120, and 240 min Torin1. Values are plotted as the mean ± SEM (≥97 cells per condition, n=3-5).

### LARP1 knockout results in global decreased mRNA stability of 5′TOP mRNAs

In addition to its role in 5′TOP translational repression, LARP1 has been reported to stabilize and protect mRNAs from degradation (17, 18, 24, 31, 32). It is currently unclear whether this protective role of LARP1 is restricted to 5′TOP mRNAs, TOP-like mRNAs, or affects all mRNAs (30). To study the effect of LARP1 and 4EBP1/2 loss on global gene expression in actively growing cells, we extracted total RNA from our cell lines and performed RNA-seq. The canonical and 5′TOP mRNA cell lines of the same genotype were sequenced together as biological replicates since expression of different reporter mRNAs should not have a global effect on gene expression and combining the independently generated cell lines reduces off-target effects caused by the CRISPR KO or shRNA KD.

To determine the effect of LARP1 loss on gene expression, we compared the transcriptome of LARP1 KO and WT cells (12,403 transcripts, CPM>1, pseudogenes excluded). As expected, LARP1 transcript levels were downregulated in the KO cell lines to 30% of WT levels (Fig. 4A, Supplementary Table 1). Volcano-plot analysis of the transcriptome changes of KO vs WT cell lines (biological replicates: n=2 for LARP1 WT, n=8 for LARP1 KO) revealed that the most significantly affected mRNAs are endogenous 5′TOP mRNAs, which are almost all downregulated in the LARP1 KO cells. Analyzing all canonical 5′TOP mRNAs (Supplementary Table 2), 70 out of 94 5′TOPs are found to be significantly down-regulated (log2 FC ≤ −0.5, -log10 p-value ≥ 5), as well as 85 significantly downregulated non-5′TOP RNAs and 40 significantly upregulated non-5′TOP RNAs. In contrast, the previously identified TOP-like mRNAs, which were predicted to be translationally regulated by LARP1 based on sequence similarity (30), were mostly unaffected in their expression.

**Fig. 4.**
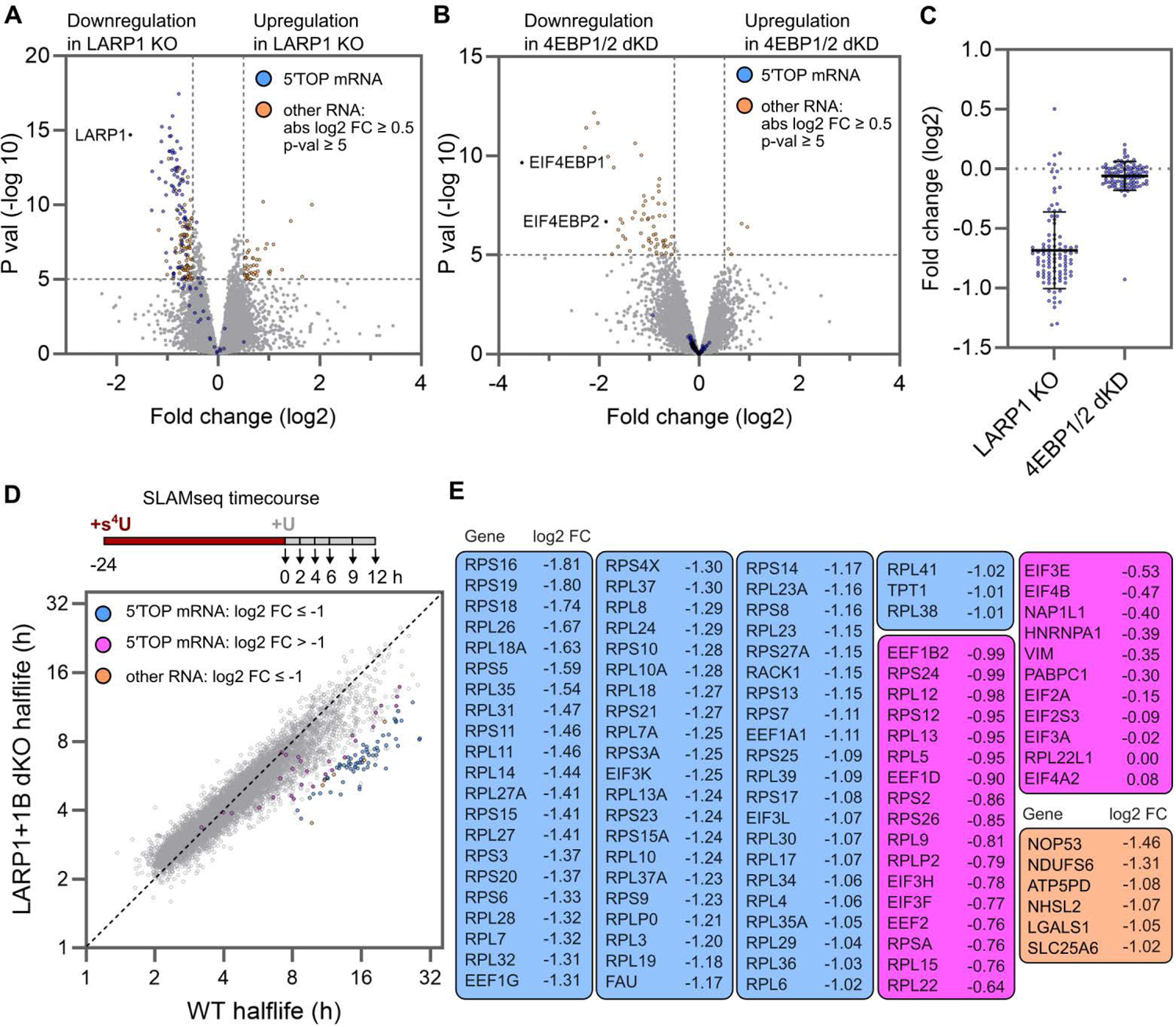
Loss of LARP1 results in selective destabilization of 5′TOP mRNAs. (**A**) Differential gene expression analysis of LARP1 KO cells compared to WT cell lines. Significantly down- and upregulated mRNAs are highlighted (-log10 p-value ≥ 5, and abs log2 FC ≥ 0.5) indicating all classical 5′TOP mRNAs (blue circles) as well as all significant non-5′TOP mRNAs (orange circles). For WT, data is the average of n=2 biological replicates (canonical and 5′TOP reporter cell lines, averaged). For LARP1 KO, data is the average of n=8 biological replicates (4 KO clones of each reporter cell line). (**B**) Differential gene expression analysis of 4EBP1/2 dKD reporter cell lines compared to WT reporter cell lines. Significantly down- and upregulated mRNAs (blue and red box, respectively) are highlighted (orange circles, -log10 p-value ≥ 5, and abs log2 FC ≥ 0.5). 5′TOP mRNAs (blue) are unaffected by 4EBP1/2 dKD. Data shown for n=2 biological replicates (canonical and 5′TOP reporter cell lines, averaged). (**C**) Fold change of 5′TOP mRNAs shown in (A) and (B). Mean fold change (log2) in LARP1 KO cells = −0.68, in 4EBP1/2 dKD cells = −0.06. (**D**) Correlation plot of mRNA half-lives in LARP1/1B dKO compared to WT reporter cells. Experimental setup of global mRNA stability analysis using s^4^U labelling (SLAM-seq) shown on top, with 24hr s^4^U labelling, wash-out, and timepoint collection. RNAs with significantly decreased stability (log2 FC ≤ −1) are highlighted for 5′TOP mRNAs (blue circles) and non-5′TOP mRNAs (orange circles), as well as non-significantly downregulated 5′TOP mRNAs (magenta circles). (**E**) List of all RNAs highlighted in (D), ranked by log2 FC for 5′TOP mRNAs and non-5′TOP mRNAs separately.

In order to determine if depletion of 4EBP1/2 also affected the level of 5′TOP transcripts, we compared the transcriptome of 4EBP1/2 dKD cells to WT cells (13,832 transcripts, CPM>1, pseudogenes excluded). Consistent with shRNA KD of 4EBP1/2, we found the levels of these two transcripts to be reduced by 91% (4EBP1) and 71% (4EBP2), and that LARP1 expression was unaltered in both cell lines. A small set of 65 transcripts showed significantly altered expression, however, these do not include canonical 5′TOP mRNAs (Fig. 4B, orange circles, abs log2 FC ≥ 0.5, -log 10 p-value ≥ 5, Supplementary Table 3). Additionally, the upregulated mRNAs did not match mRNAs described to be sensitive to eIF4E levels in mice (33). In contrast to the role of 4EBP1/2 in regulating translation during mTOR inhibition, these results indicate that their loss does not alter the levels of 5′TOP transcripts (Fig. 4C).

The selective downregulation of endogenous 5′TOP mRNAs we observed in the absence of LARP1 suggested that LARP1 specifically stabilizes 5′TOP mRNAs. To confirm that the changes in steady-state expression were caused by mRNA destabilization and were not due to changes in transcription, we performed global mRNA half-life measurements using metabolic 4-thiouridine labelling (SLAM-seq (16)). WT and LARP1/1B dKO cells of both canonical and 5′TOP cell lines were incubated with 4-thiouridine for 24 hours, followed by wash-out and harvesting of cells over a 12-hour time course. Half-lives of mRNAs were calculated using a single-exponential decay fit for 9,837 transcripts (R^2^ ≥ 0.75, pseudogenes excluded, Supplementary Table 4). In agreement with previous measurements of mammalian mRNA half-lives, the global median half-life for both WT and LARP1/1B dKO HeLa cell lines was ∼4 hours (Fig. S11A), indicating that loss of LARP1 does not globally destabilize all mRNAs. Correlation analyses showed a high correlation in mRNA half-lives among the four cell lines (r=0.90-0.94), allowing us to compare the mRNA half-lives in WT vs LARP1/1B KO cell lines (Fig. 4D, n=2). In agreement with our RNA-seq results, nearly all 94 5′TOP mRNAs detected in the SLAM-seq experiment have decreased mRNA stability, with 66 5′TOP mRNAs changing by > 2-fold (Fig. 4D). Some 5′TOP mRNAs seem largely unaffected by LARP1 loss (e.g. PABPC1 and EIF3A), suggesting the potential involvement of additional stabilizing factors. Furthermore, the length of the 5′TOP motif or presence of a PRTE motif within the 5′UTR does not correlate with the change in mRNA half-lives (Fig. S11B). Only six non-5′TOP mRNAs were found to be destabilized > 2-fold and three of these transcripts (NOP53, LGALS1, SLC25A6) have annotated transcription start sites that contain 5′ TOP motifs suggesting that they could be similarly regulated in HeLa cells. Taken as a whole, our results support a model of LARP1-mediated stabilization that is highly selective for 5′TOP mRNAs.

## Discussion

In this study, we employed single-molecule imaging to study the regulation of translation of 5′TOP mRNAs upon mTOR inhibition that allowed us to directly quantify the effect of LARP1 and 4EBP1/2. By imaging and quantifying the translation status of individual mRNAs, we find that 4EBP1/2 plays a dominant role compared to LARP1 in mediating 5′TOP translational repression under short-term (30min-4h) pharmacological inhibition of mTOR using Torin1 in HeLa cells. Previously, studies that used genome-wide ribosome or polysome profiling determined that LARP1 and 4EBP1/2 regulate 5′TOP translation during mTOR inhibition (3, 4, 10, 11, 30, 34, 35), however, the magnitude of their respective contributions was difficult to measure due to the inherent limitations of these approaches. We believe this highlights the power of single-molecule imaging methods for quantifying translation in livings cells in order to determine the specific effects of translation factors.

While our results indicate that 4EBP1/2 is the critical factor in mediating 5′TOP translational repression, the underlying molecular mechanism is not entirely clear. Although we cannot exclude the possibility of a still unknown factor acting downstream of 4EBP1/2, we favor a model where the translation of 5ʹ TOP mRNAs is more sensitive to active eIF4E levels. *In vitro* experiments have determined that eIF4E binds with ∼3-fold weaker affinity to m^7^GTP-capped oligonucleotides with a +1 cytosine than either purine, which is consistent with translation of 5′TOP mRNAs being slightly worse than a non-5′TOP mRNA when mTOR is active and then preferentially repressed when available eIF4E levels become extremely limited during mTOR inhibition (14, 36, 37). This model is also in line with previous work that found inducible-overexpression of eIF4E to specifically upregulate the translation of 5′TOP mRNAs (38), as well as recent work that showed that 5′TOP mRNAs are less sensitive to mTOR inhibition in acutely PABPC1-depleted cells where global mRNA levels are reduced (Marc Fabian, personal communication (39)).

Interestingly, in the X-ray structure of human eIF4E in complex with m^7^pppA, the C-terminal tail of eIF4E adopts a conformation that enables Thr205 to form a hydrogen bond with the exocyclic amino of the adenine base (40). While the position of the eIF4E C-terminal tail has not been determined when bound to longer RNA sequences, phosphorylation of Ser209 is known to enhance translation indicating that additional residues in this region may have functional roles (41). Alternatively, other canonical translation factors (e.g. eIF4G or 4EBP1/2) may also contribute to 5′TOP specificity through additional interactions (42, 43).

While LARP1 may not be the key repressor in 5′TOP translational regulation, our data supports a major role of LARP1 in mediating 5′TOP mRNA stability when mTOR is active. Previous work established a link between LARP1 and mRNA stability, with LARP1 binding both PABP and the polyA tail and inhibiting deadenylation (17-19, 21, 31, 32, 44, 45). It has been unclear whether this protective role is restricted to 5′TOP mRNAs as binding to PABP/polyA is anticipated to not be selective, and crosslinking studies have found LARP1 complexed with thousands of mRNAs (21–23). Our results show a highly selective destabilization of nearly all 5′TOP mRNAs upon loss of LARP1, with virtually all other mRNAs being largely unaffected. Similarly, a recent study found that loss of LARP1 resulted in rapid deadenylation of short polyA tails of all mRNAs, with 5ʹTOP mRNAs being more affected than other types of mRNAs (24). It is possible that differences in LARP1 depletion or measurement of mRNA stability or polyA-tail length may account for the differences in specificity for 5ʹTOP mRNAs between the studies.

Previous studies have focused on the role of LARP1 in protecting 5′TOP mRNAs in mTOR inhibited cells, as LARP1 has been shown to be recruited to 5′TOP mRNAs upon mTOR inhibition (21, 44). Our findings raise the intriguing question of how LARP1 can be specifically recruited to 5′TOP mRNAs when mTOR is active. While it has been proposed that LARP1 can interact with its La-motif with both the 5′TOP motif and PABP (32), it is not clear that this interaction is compatible with eIF4F or translation. Recent structural work of the human 48S preinitiation complex suggests that there could be a “blind spot” of ∼30nts adjacent to the cap that might allow LARP1 to bind the 5ʹTOP sequence (46). Importantly, we do not observe any correlation between change in mRNA half-lives with either length of 5′TOP motif or presence of other pyrimidine-rich elements (PRTE) in the 5ʹUTR suggesting that the position of the pyrimidines directly adjacent to the cap is necessary for this effect on mRNA stability. Interestingly, LARP1 was shown to promote the localization of ribosomal mRNAs in a PRTE-dependent manner but did not require the more strict 5′ TOP motif suggesting that LARP1’s interaction with ribosomal mRNAs and its functional consequence could be context-dependent (47).

While our translation site imaging experiments are limited to the characterization of two reporter mRNAs (5′TOP and non-5′TOP), we have shown that the results are consistent with previous studies that characterized endogenous ribosomal protein mRNAs, however, single-molecule experiments in living cells allow more accurate quantification of the effect of loss of LARP1 and 4EBP1/2. We anticipate that similar results would be obtained with 5′TOP sequences derived from ribosomal protein mRNAs other than RPL32, though the magnitude of the difference in translation repression could be different if compared to another non-5′TOP transcript. Additionally, the continued development of methodologies for imaging translation of single mRNAs for extended time periods and the interplay of translation with mRNA decay will enable the dynamics of mTOR regulation to be quantified in greater detail (48, 49).

## Materials and Methods

### Materials

All antibodies used in this study are listed in Table 1. All chemicals, plasmids, viruses, cell lines, sgRNA, and shRNA used in this study are listed in Supplementary Table 5.

**Table 1.** Antibodies.

| Primary Antibodies | Provider | Cat. # |
| --- | --- | --- |
| LARP1 | Bethyl Labs | A302-087A |
| LARP1 | Cell Signaling Technology | 70180 |
| α-Tubulin | Cell Signaling Technology | 3873 |
| mTOR | Cell Signaling Technology | 2983 |
| Phospho-RPS6 | Cell Signaling Technology | 2211 |
| Phospho-4E-BP1 (Ser65) | Cell Signaling Technology | 9451 |
| 4E-BP1 | Cell Signaling Technology | 9452 |
| 4E-BP2 | Cell Signaling Technology | 2845 |
| **Secondary Antibodies** |  |  |
| IRDye® 680RD Goat anti-Mouse IgG | LI-COR | 926-68070 |
| IRDye® 800CW Goat anti-Rabbit IgG | LI-COR | 926-32211 |

### Cell culture

The HeLa-11ht cell lines expressing either RPL32 5′TOP SunTag or non-5′TOP canonical SunTag mRNAs used in this study were previously generated in the Chao lab (Wilbertz et al. 2019), and the corresponding plasmids are available from Addgene (#119946, #119945). The reporter cell lines were grown in 10% FCS-DMEM medium containing 4.5 g/L glucose, 100 µg/ml penicillin and streptomycin, 4mM L-glutamine, and 10% fetal bovine serum (FBS) at 37°C and 5% CO2. To maintain the reverse tetracycline-controlled transactivator (rtTA2-M2) for inducible expression, the medium was supplemented with 0.2 mg/ml G418.

HEK293T cells used for lentivirus production were grown in 10% FCS-DMEM medium containing 4.5 g/L glucose, 100 µg/ml penicillin and streptomycin, 4mM L-glutamine, and 10% fetal bovine serum (FBS) at 37°C and 5% CO2.

### Validation of transcription start sites for 5′TOP and canonical SunTag transcripts

For mapping of transcription start sites, total RNA was converted into full-length adapterligated double-stranded cDNA using the TeloPrime Full-Length cDNA Amplification Kit V2 (Lexogen), which employs a cap-specific adapter selective for intact mRNAs. cDNA 5′-terminal sequences were amplified by PCR using a gene-specific primer and TeloPrime forward primer, cloned into the pCR-Blunt vector (Thermo Fisher Scientific), and sequenced.

### CRISPR knockout cell line generation

To generate LARP1 CRISPR knockout clones, parental HeLa-11ht cell lines expressing the reporter mRNAs were transiently co-transfected with two Cas9 plasmids, each containing Cas9 and a sgRNA targeting a sequence within exon 4 of LARP1, enhancing efficiency of knockout cell line generation (50, 51). Transient transfections were performed following the manufacturer’s instructions using Lipofectamine 2000 (Invitrogen) and Opti-MEM (Thermo Fisher Scientific). Two days after transient transfections, highly transfected cells were single-cell sorted into 96-well plates for monoclonal selection (10% highest mCherry positive cells, using Cas9-T2A-mCherry). Single clonal cell populations were screened for loss of LARP1 by immunostaining (Bethyl Labs #A302-087A) in 96-well plates. For both 5′TOP- and canonical-SunTag cell lines, four knockout clones each were verified by western blot for loss of LARP1 protein expression.

To generate LARP1B CRISPR knockout clones, the LARP1 knockout reporter cell lines were similarly co-transfected with two Cas9 plasmids carrying two different sgRNAs targeting sequences within exon 4 of LARP1B. Following the same steps as described for generating LARP1 knockout cell lines, clonal cell populations were screened by PCR for presence of truncated LARP1B alleles, and subsequently verified by genomic DNA sequencing and cDNA amplification.

### shRNA stable knockdown cell lines

To generate 4EBP1/2 double KD cells, two lentiviruses expressing different resistance genes were used that contained shRNA sequences from the RNAi Consortium public library that were previously described (Thoreen et al. 2012). The 4EBP1 shRNA lentivirus carrying puromycin resistance was purchased as lentiviral particles (Sigma-Aldrich). The 4EBP2 shRNA was cloned into the pLKO.1_BlastR lentiviral backbone (52). To produce 4EBP2 shRNA lentivirus, HEK293T cells were co-transfected with the 4EBP2 shRNA, the psPax2 envelope and the vsv-G packaging plasmids using Fugene6 (Promega) according to the manufacturer’s instructions. Supernatant containing viral particles was harvested daily for the next four days, centrifuged, and filtered through a 0.45 μM filter to remove cell debris.

The viral particles were concentrated by precipitation using the Lenti-X concentrator (Clontech) and resuspended in cell culture medium. For infection of HeLa cells expressing the reporter mRNAs, 10,000 cells were seeded in 12-well dishes and co-infected the next day with 4EBP1 and 4EBP2 shRNA viruses in medium supplemented with 4 µg/ml polybrene (Merck). Cells were grown until confluency and reseeded into 6-well dishes prior to addition of 1 µg/ml puromycin (InvivoGen) and 5 µg/ml blasticidin (InvivoGen). Uninfected HeLa-11ht cells were used to determine the minimal antibiotic concentrations that result in lethality within 2-5 days. Double resistant cell lines with dual integration of 4EBP1/2 shRNAs were validated by western blot (Cell Signaling Technology #9452, #2845) for efficient stable knockdown of the targeted proteins.

### Genomic DNA extraction

For genotyping of CRISPR edited cell lines, cells were harvested by trypsinization, and the genomic DNA was extracted using the DNeasy kit (Qiagen) according to the manufacturer’s instructions. Primers specific to the target genes were designed using the Primer-Blast tool (https://www.ncbi.nlm.nih.gov/tools/primer-blast/). Genomic DNA was amplified using Phusion High-Fidelity polymerase (NEB), PCR products were cleaned using a PCR purification kit (Qiagen), and purified PCR products were cloned into the pCR-Blunt vector (Thermo Fisher Scientific). For each cell line, a minimum of 10 clones were isolated and analyzed by Sanger sequencing to identify all edited alleles.

### SDS-PAGE and Western Blotting

For protein extraction, cells were harvested by trypsinization and lysed in RIPA buffer (150 mM NaCl, 50 mM Tris, 0.1% SDS, 0.5% sodium deoxycholate, 1% Triton X-100) supplemented with 1x protease inhibitor (Bimake.com) and SuperNuclease (Sino Biological). Cell lysate was centrifuged at 12,000 rpm for 10 minutes to remove cell debris, and the supernatant was loaded on a 4-15% tris-glycine gel (BioRad) using 1x Laemmli buffer (BioRad) supplemented with 100 mM DTT (VWR). Following SDS-PAGE, proteins were transferred onto nitrocellulose or PVDF membranes by semi-dry transfer (Trans-Blot Turbo) and blocked in 5% BSA-TBST buffer (TBS supplemented with 0.1% Tween-20) for 1hr at RT. Primary antibodies were incubated overnight at 4°C in TBST or Intercept® blocking buffer + 0.1% Tween-20. The next day, the membrane was washed 3-5 times in TBST, and incubated for 1hr at RT with the fluorescent secondary antibodies diluted 1:10,000 in LI-COR blocking buffer + 0.1% Tween-20 (supplemented with 0.01% SDS for PVDF membranes). Following 3-5 washes in TBST, membranes were transferred to PBS and antibody fluorescence was detected at 700 and 800 nm using an Odyssey infrared imaging system (LI-COR).

### Total RNA extraction and cDNA synthesis

For total RNA extraction, cells were harvested by trypsinization and lysed in RNA lysis buffer following the RNA Miniprep kit (Agilent). Genomic DNA contamination was reduced by on-column DNase digestion as described in the manual, and purified RNA was stored at −80°C. For validation of LARP1B CRISPR knockout, total RNA was reverse transcribed to cDNA, which was used as the template for cDNA amplification of edited LARP1B transcripts. LARP1B transcripts were amplified as described above for gDNA validation.

### Live-cell imaging

For live-cell imaging, cells were seeded at low density (20-30,000) in 35mm glass-bottom µ-dishes (Ibidi) and grown for 2-3 days. On the day of imaging, the cells were incubated with JF549 or JF646 red/far-red dyes (HHMI Janelia Research Campus) to label the MCP-Halo coat protein for 30min, unbound dye was removed (3 washes, PBS), and cells were kept in culture medium until imaging. For induction of reporter mRNAs, 1 µg/ml doxycycline (Sigma-Aldrich) was added to each dish at appropriate time points before each imaging session (30 minutes) to ensure the same duration of doxycycline induction at the start of imaging for all dishes of an experiment.

At the start of imaging of each dish, culture medium was replaced with FluoroBrite imaging DMEM (Thermo Fisher Scientific) supplemented with 10% FCS, 2 mM glutamine, and 1 µg/ml doxycycline. To inhibit mTOR, cells were treated with 250 nM Torin1 at various timepoints, which was maintained throughout the imaging session. To inhibit translation, cells were treated with 100 μg/ml puromycin 5 min prior to the start of imaging, which was maintained throughout imaging. To deplete elongating ribosomes, cells were treated with 3 µg/ml harringtonine 10 min prior to the start of imaging, which was maintained throughout imaging. For all experiments, the start of the 30 min imaging window was recorded as the timepoint shown in the figures (e.g. imaging 60-90 min after Torin1 addition = 60 min timepoint).

Cells were kept at 37°C and 5% CO2 throughout image acquisition. All dual-color live-cell imaging was performed on an inverted Ti2-E Eclipse (Nikon) microscope equipped with a CSU-W1 scan head (Yokogawa), two back-illuminated EMCCD cameras iXon-Ultra-888 (Andor) with chroma ET525/50 m and ET575lp emission filters, and an MS-2000 motorized stage (Applied Scientific Instrumentation). Cells were illuminated with 561 Cobolt Jive (Cobolt), 488 iBeam Smart, 639 iBeam Smart (Toptica Photonics) lasers, and a VS-Homogenizer (Visitron Systems GmbH). Using a CFI Plan Apochromat Lambda 100x Oil/1.45 objective (Nikon), images were obtained with a pixel size of 0.134 μm. To allow for simultaneous tracking of mRNA and translation sites, both channels were simultaneously acquired by both cameras at 20 Hz for 100 frames in a single Z-plane (5 sec movies).

### Live-cell data analysis

For image analysis, the first 5-14 frames of each movie (500ms) were selected for single particle tracking. First, images were corrected for any offset between the two cameras using TetraSpeck fluorescent beads acquired on each imaging day. Using the FIJI (53, 54) descriptor-based registration plugin (55) in affine transformation mode, a transformation model was obtained to correct the bead offset, and applied to all images of an imaging day using a custom macro (56). Subsequently, fine correction of remaining offsets between images were corrected for each dish individually using the FIJI translate function run in a custom macro, correcting for offsets occurring progressively throughout an imaging session.

Single-particle tracking and translation site quantification was performed as described previously (56). In short, using the KNIME analytics platform and a custom-build data processing workflow, regions-of-interest (ROIs) were manually annotated in the mRNA channel, selecting cytosolic regions with multiple bright spots. Importantly, annotation solely in the mRNA channel excludes any bias in selection attributable to the translational state of the cell. Next, spots in the ROIs corresponding to single mRNAs were tracked using Track-Mate (57) integrated in KNIME, using the “Laplacian of Gaussian” detector with an estimated spot radius of 200 nm and sub-pixel localization. Detection thresholds were adjusted based on the SNR of images and varied between 1.25-2. For particle-linking, the parameters linking max distance (600 nm), gap closing max distance (1200 nm), and gap closing max frame gap (2) were optimized for single-particle tracking of mRNAs. To assay whether an mRNA is translating, the mean intensity of the SunTag channel was measured at the coordinates of each mRNA spot, and quantified as fold-change / ROI background intensity. A cut-off of <1.5 fold/background was determined to classify an mRNA as non-translating based on calibration data using the translation inhibitor harringtonine.

For data visualization, the fraction of translating mRNAs per cell and translation site intensities were plotted using SuperPlots (https://huygens.science.uva.nl/SuperPlotsOfData/), showing all data points together with the mean (± SD) of each biological replicate (58).

### RNA-seq

For RNA-seq, total RNA samples extracted using the RNA Miniprep kit (Agilent), were assessed for RNA integrity using the Agilent Tapestation, and library preparation was performed using the Illumina TruSeq Stranded mRNA reagents according to the manufacturer’s protocol. Libraries were sequenced on the Illumina HiSeq2500 (GEO submission In progress: single reads, 50 cycles) or NovaSeq6000 platforms (GEO submission In progress: paired-end reads, 100 cycles).

For our analysis, we used a reference list of experimentally validated 5′TOP mRNAs (30), expanded with additional experimentally verified 5′TOP mRNAs (RACK1, EIF3K) as well as computationally predicted 5′TOP mRNAs with known roles in translation (EIF2A, EIF2S3, EIF3L, EIF4A2, and RPL22L1) (30). The presence or absence of a PRTE in the 5′UTR was taken from Hsieh et al. 2012 (59), and expanded by manual annotation for the subset of 5′TOP mRNAs not listed (Supplementary Table 2).

Sequenced reads were aligned against the human genome (GENCODE GRCh38 primary assembly, https://www.gencodegenes.org/human/release_38.html) using R version 4.1.1 with Bioconductor version 3.13 to execute the qAlign tool (QuasR package, version 1.32.0, (60)), with default parameters except for aligner = “Rhisat2”, splicedAlignment = “TRUE”, allowing only uniquely mapping reads. Raw gene counts were obtained using the qCount tool (QuasR) with a TxDb generated from gencode.v38.primary_assembly.annotation.gtf as query, with default parameters counting only alignments on the opposite strand as the query region. The count table was filtered to only keep genes which had at least 1 cpm in at least 3 samples.

### SLAM-seq

For metabolic labelling (SLAMseq), HeLa-11ht cell lines were incubated with a dilution series of 4-thiouridine (S4U) for 24h, exchanging S4U-containing media every 3h according to the manufacturer’s instructions. S4U cytotoxicity was assessed using a luminescent cell viability assay, and the half-maximal inhibitory concentration (IC50) was calculated at 209 μM (n=2). Based on the IC50, a trial RNAseq was conducted with 24hr S4U labelling using a dilution serios (0, 3, 6, 12, 24 and 48μM) of S4U and exchanging media every 3hr. The 12 μM S4U concentration was selected as the optimal experimental S4U concentration with minimal effects on gene expression for all cell lines.

For assessing global RNA half-lives, HeLa-11ht cell lines were labelled with 12 μM S4U for 24hr (exchanging media every 3h), labelling was stopped using 100x excess uridine (1.2 mM), and cells were harvested at timepoints 0, 2, 4, 6, 9, and 12hr after the uridine quench. For isolation of total RNA, RNA was extracted using an RNA miniprep kit (Agilent) with on-column DNase digestion, followed by iodoacetamide treatment and EtOH precipitation of modified total RNA. For RNA-seq, total RNA was assessed for RNA integrity using the Agilent Tapestation, and library preparation was performed using the Illumina TruSeq Stranded Total RNA Library Prep Gold kit according to the manufacturer’s protocol. Libraries were sequenced on the Illumina NextSeq500 (GEO submission In progress: single reads, 75cycles). Samples were submitted as three independent replicates (cells harvested on separate days), with the exception of one sample with only two replicates (one sample lost during sequencing).

In total ∼3.7 billion SLAMseq reads were produced corresponding to ∼52M reads per replicate. 4-thiouridine incorporation events were analyzed using the SlamDunk software (v0.3.4) for SLAMseq analysis (61). SLAMseq reads were first reverse complemented to match the hard-coded assumed SlamDunk orientation using fastx_reverse_complement from the FASTX-toolkit (http://hannonlab.cshl.edu/fastx_toolkit) with default settings. The resulting fastq files were mapped to the reference genome (GENCODE GRCh38 primary assembly) with slamdunk map and parameters −5 0 -ss q. The mapped reads were subsequently filtered to only retain intragenic mappings according to the reference transcriptome (GRCh38, GENCODE v33) with a high identity using slamdunk filter and parameters -mi 0.9. SNP variants in the samples were called with slamdunk snp with parameters -c 1 -f 0.2. The SNP variants from all samples were combined in a single master vcf file by indexing the individual vcf file using tabix of the htslib package (https://github.com/samtools/htslib) and merging using vcf-merge from the VCFtools package (62). 4-thiouridine incorporation and conversion rates were calculated separately for exonic gene segments of the reference transcriptome using slamdunk count with default parameters and the master SNP vcf file for SNP filtering. The exonic segments counts were then aggregated to obtain gene level total mapping reads, multimapping reads and converted reads counts. During aggregation, total and converted counts from exonic segment with multimappers were downweighted by the fraction of mutimappers over the total mapped reads of the the exonic segment (fm). Finally, gene conversion rates were calculated as the number of the gene-level aggregated converted reads over the gene-level aggregated total read counts.

Gene conversion rates in each context were fitted to an exponential time decay model to obtain gene half-life estimates. Fitting was performed by non-linear least squares using the R stats::nls function. Example fit command: *fit <-nls(rates ∼ exp(a+k*timepoints),control=list(minFactor=1e-7, tol=1e-05,maxiter = 256))* where rates are the conversion rates for the gene (including all replicates) and timepoints are the corresponding times in hours. The half-life (in min) was obtained from the fitted coefficient (t1/2= −1/k*ln(2)*60). Half-life estimates and the fitting R^2^ values are listed in Suppl. Table 4 for all transcripts with R^2^ ≥ 0.75 in all conditions.

### Statistical Analysis

For live-imaging, biological replicates (n) were defined as independent days of imaging. Statistical analyses were performed using GraphPad Prism, with n numbers and statistical tests described in the figure legends. Technical replicates within biological replicates were pooled before statistical tests.

For RNA-seq analysis of LARP1 WT cells, biological replicates (n) were defined as independent clonal cell lines (canonical and 5′TOP mRNA cell lines, n=2). For RNA-seq of LARP1 knockout clones, four single-cell derived clones were sequenced for both canonical and 5′TOP cell lines and treated as independent biological replicates (n=8 total). For each n, three independent replicates (cells harvested on separate days) were submitted for RNA-seq and averaged before statistical tests. Differential gene expression was calculated with the Bioconductor package edgeR (version 3.34.0, (63)) using the quasi-likelihood F-test after applying the calcNormFactors function, obtaining the dispersion estimates and fitting the negative binomial generalized linear models.

For SLAM-seq analysis, biological replicates (n) were defined as independent clonal cell lines for both LARP1 WT and LARP1/1B dKO cells (canonical and 5′TOP mRNA cell lines, n=2). For analysis of changes in mRNA stability, the estimated half-lives were averaged for n1 and n2, and changes in mRNA stability were calculated between genotypes (log2 FC). Significant differences in mRNA stability were classified with abs log2 FC ≥ 1, excluding spurious transcripts overlapping in sequence with known 5′TOP mRNAs (read-through transcripts AC135178.3, AP002990.1, AC245033.1, lncRNA AL022311.1, MIR4426, MIR3654).

## Acknowledgments

We thank Veronika Herzog for her advice on SLAMseq methodology, S. Smallwood and H.-R. Hotz for support with transcriptomics, F. Voigt, L. Gelman, L. Plantard, and S. Reither for microscopy and image analysis support, H. Kohler for cell sorting and all members of the Chao lab for their input and helpful discussions. A donation was made to the Henrietta Lacks Foundation to acknowledge the use of HeLa cells in our research.

## Author contributions

T.H. performed experiments and analyzed data. P.P. and E.P. performed the SLAMseq analysis. T.H. and J.A.C. wrote the manuscript.

## Data and materials availability

All data are available in the main text or the supplementary materials. All reagents generated in this study are available upon request from the lead contact, Dr. Jeffrey A. Chao with a completed materials transfer agreement. Microscopy data in this paper will be shared by the lead contact upon request. All original code (KNIME workflows and ImageJ macros) has been deposited at Zenodo and are publicly available. Any additional information required to reanalyze the data reported in this paper is available from the lead contact upon request.

---

**Fig. S1.**
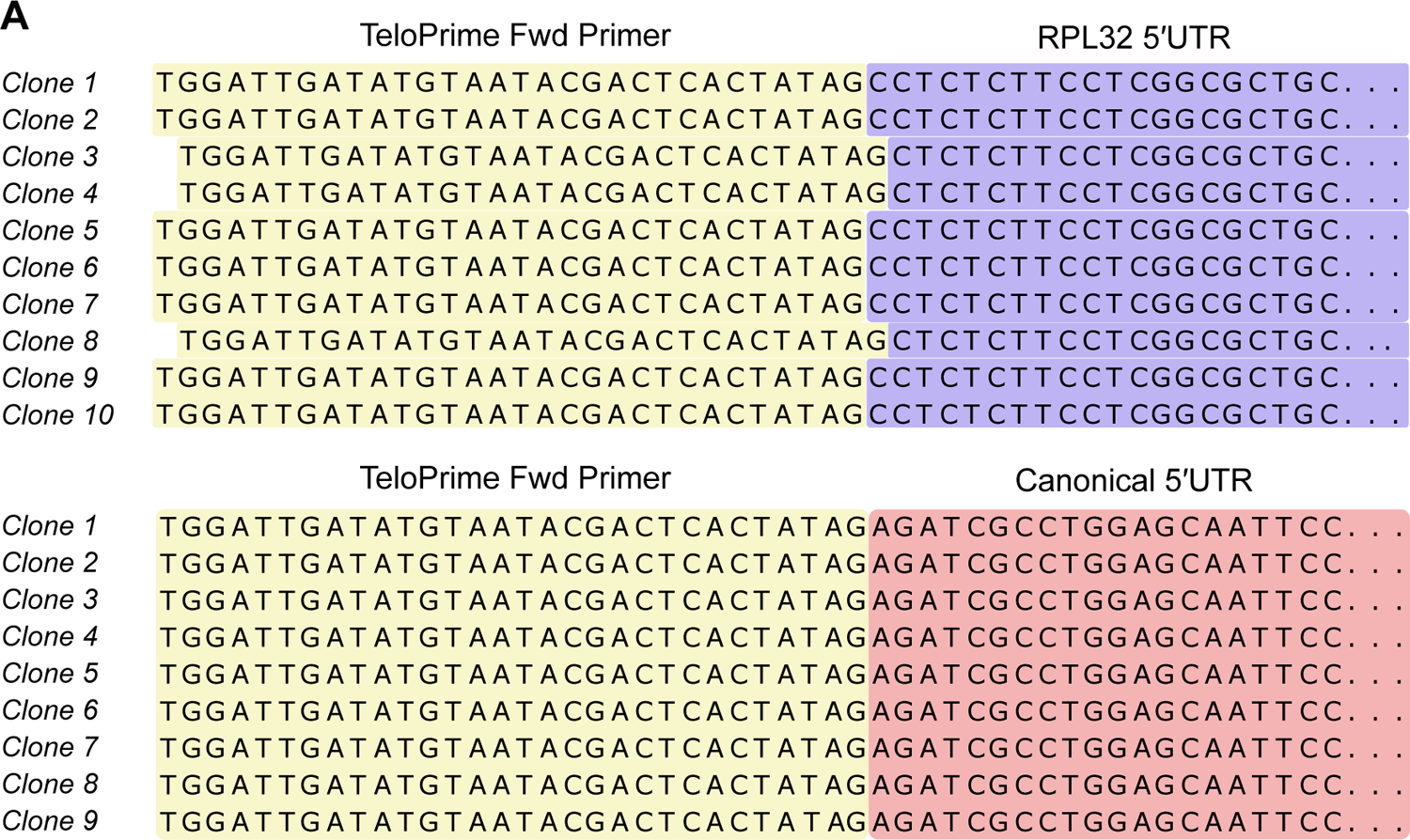
Validation of transcription start site selection. Mapping of canonical and 5′TOP reporter transcription start sites (TSS) using cap-specific adaptor ligation. Sequencing results for single clones of each reporter are shown with the TeloPrime adaptor sequence joined to the start of the reporter 5′UTR. For the 5′TOP reporter, there is some variability in the precise cytosine in the +1 position, however, all sequenced clones initiate with a cytosine and contain a 5′TOP motif.

**Fig. S2.**
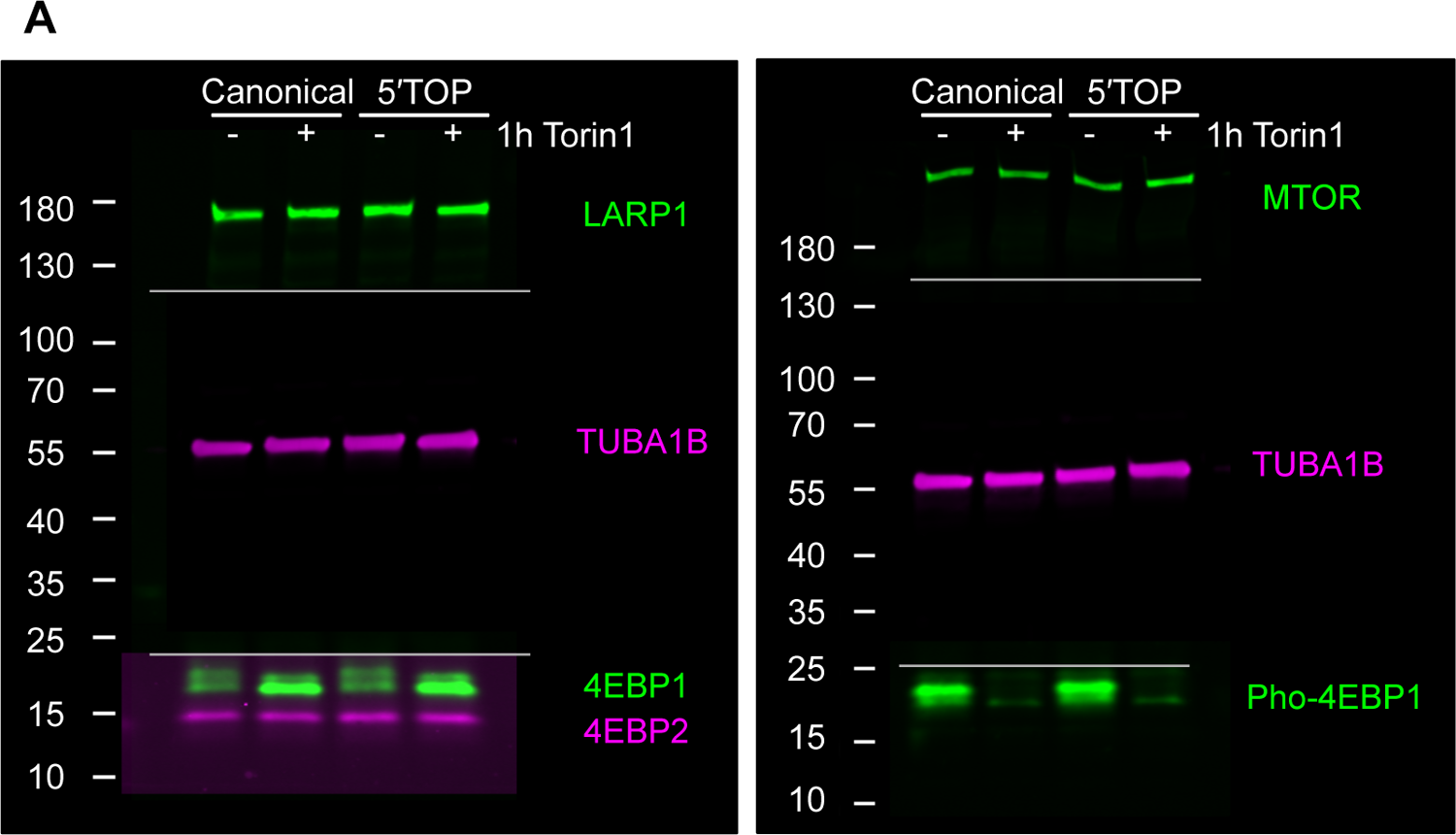
mTORC1 signaling in canonical and 5′TOP mRNA cell lines treated with Torin1. Western blot analysis of mTORC1 signaling using a phosphorylation-specific antibody against 4EBP1. Upon Torin1 addition (250nM, 1h), 4EBP1 is dephosphorylated, as seen by the lower migration size of 4EBP1 (4EBP1 antibody, left), and the disappearance of Phospho-4EBP1 (Phospho-Ser65-4EBP1 antibody, right). Lines indicate cut membrane pieces probed with different mouse (magenta) and rabbit (green) antibodies, and imaged together using two-color fluorescent imaging. Brightness and contrast were individually adjusted for each antibody shown.

**Fig. S3.**
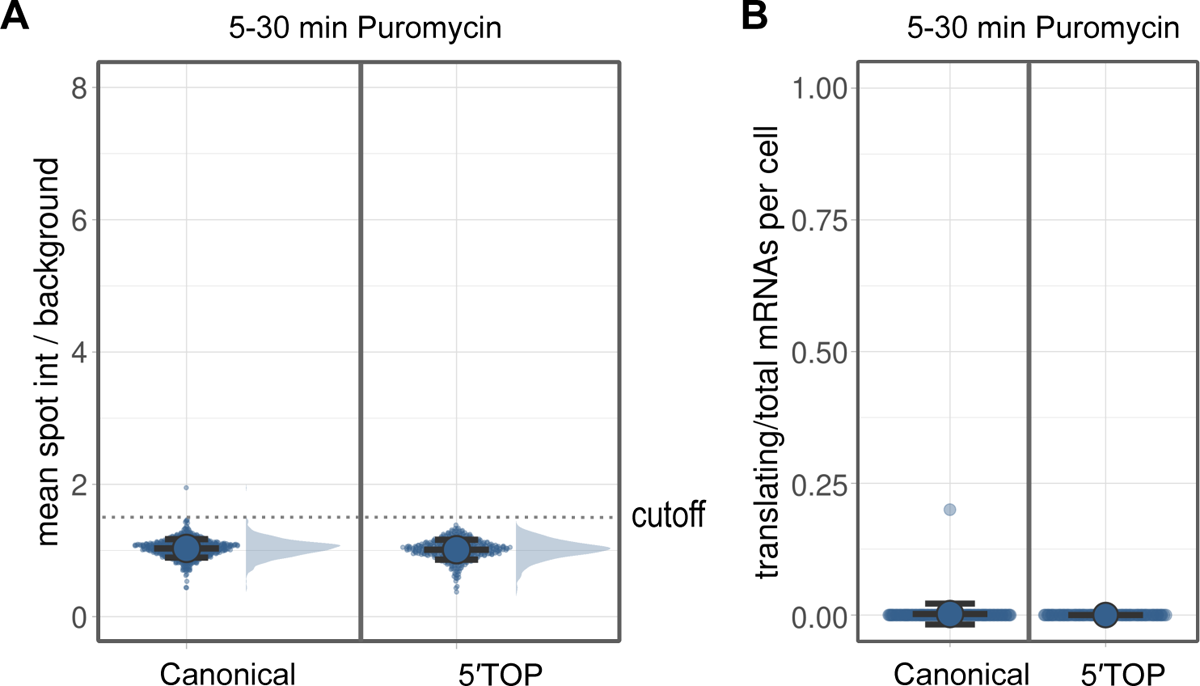
Translation site intensities following global translation inhibition. (**A**) Quantification of translation site intensities after puromycin treatment (100 μg/ml, 5-30min). SunTag intensities are plotted for >400 mRNAs per condition overlaid with the mean ± SD. A cutoff of 1.5 (dotted line) was determined to distinguish translating from non-translating mRNAs. (**B**) Fraction of mRNAs undergoing translation after puromycin treatment. Values are plotted for each cell over-laid with the mean ± SD (≥87 cells per condition).

**Fig. S4.**
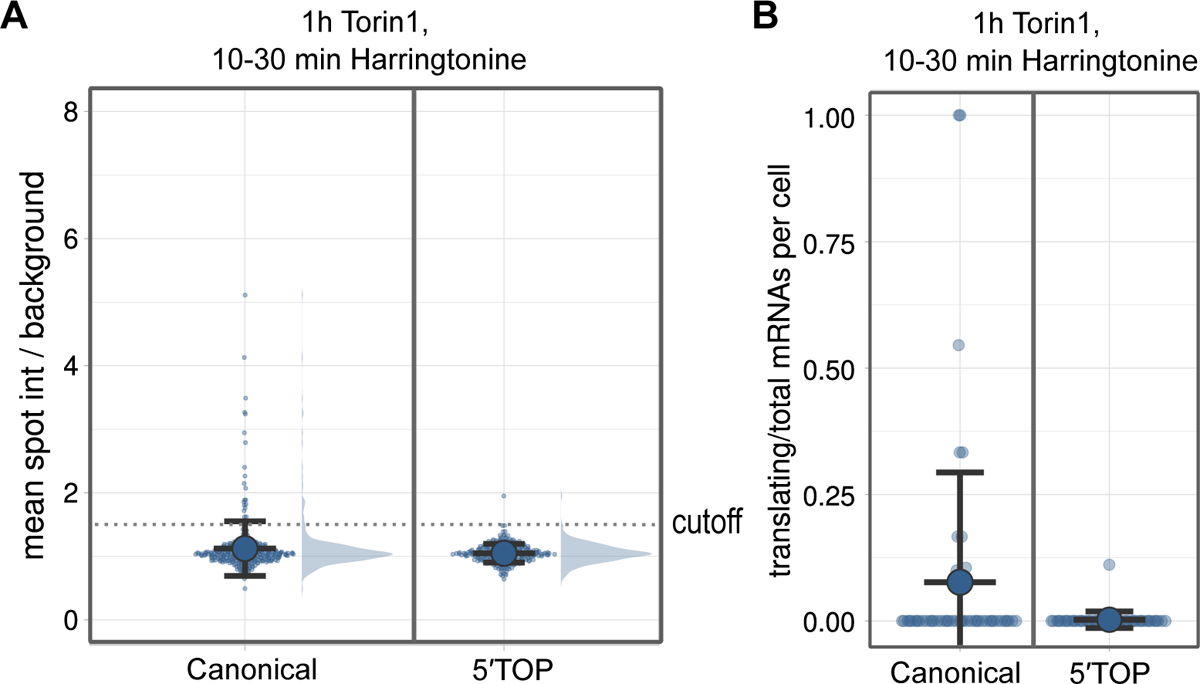
Translation site intensities following ribosome run-off. (**A**) Quantification of translation site intensities after mTOR inhibition (250nM Torin1, 1h) followed by translation inhibition (Harringtonine 3 μg/ml, 10-30min). SunTag intensities are plotted for >250 mRNAs per condition overlaid with the mean ± SD. (**B**) Fraction of mRNAs undergoing translation after mTOR inhibition (250nM Torin1, 1h) followed by translation inhibition (Harringtonine 3 μg/ml, 10-30min). Values are plotted for each cell overlaid with the mean ± SD (≥46 cells per condition).

**Fig. S5.**
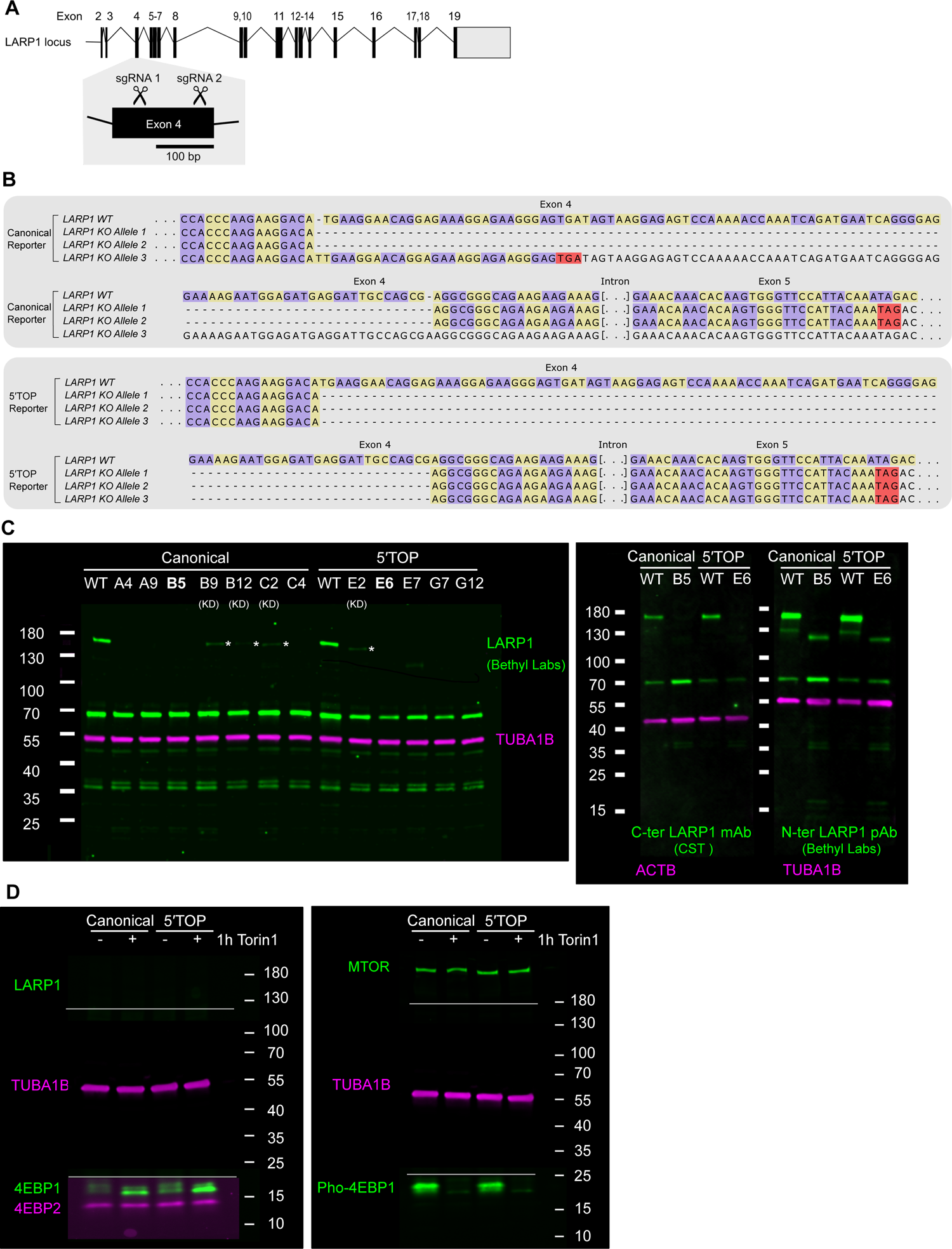
Validation of LARP1 knockout cell lines. (**A**) CRISPR-Cas9 editing strategy for deletion of LARP1 protein expression in canonical and 5′TOP mRNA cell lines. Two sgRNAs targeting exon 4 upstream of any domains of known function were used to ensure high editing efficiency in polyploidal HeLa genome (sgRNA 1 taken from (5)). (**B**) Genotyping of canonical and 5′TOP knockout (KO) clones used for single-molecule imaging (Fig. 2A, 2B, 2E), showing ∼100 bp truncations of LARP1 exon 4 resulting in frameshifting and premature termination codons. (**C**) Western Blot analysis of LARP1 protein expression in wildtype cell lines (WT) and LARP1 CRISPR-Cas9 edited single clonal cell lines. Clones B5 and E6 were used for single-molecule imaging of the canonical and 5′TOP mRNAs respectively. Note the presence of a weak shorter band of LARP1 (highlighted with *) for some knockout clones, indicating the presence of an in-frame truncated allele of LARP1 in these clones (KD). (**D**) Western blot analysis of mTORC1 signaling in canonical and 5′TOP LARP1 KO cell lines used for single-molecule imaging. Upon 1h Torin1, 4EBP1 is shifted to its dephosphorylated state, as seen by the lower migration size of 4EBP1 (4EBP1 antibody, left), and the disappearance of Phospho-4EBP1 (Phospho-Ser65-4EBP1 antibody, right). Lines indicate cut membrane pieces probed with different mouse (magenta) and rabbit (green) antibodies, and imaged together using two-color fluorescent imaging. Brightness and contrast were individually adjusted for each antibody shown.

**Fig. S6.**
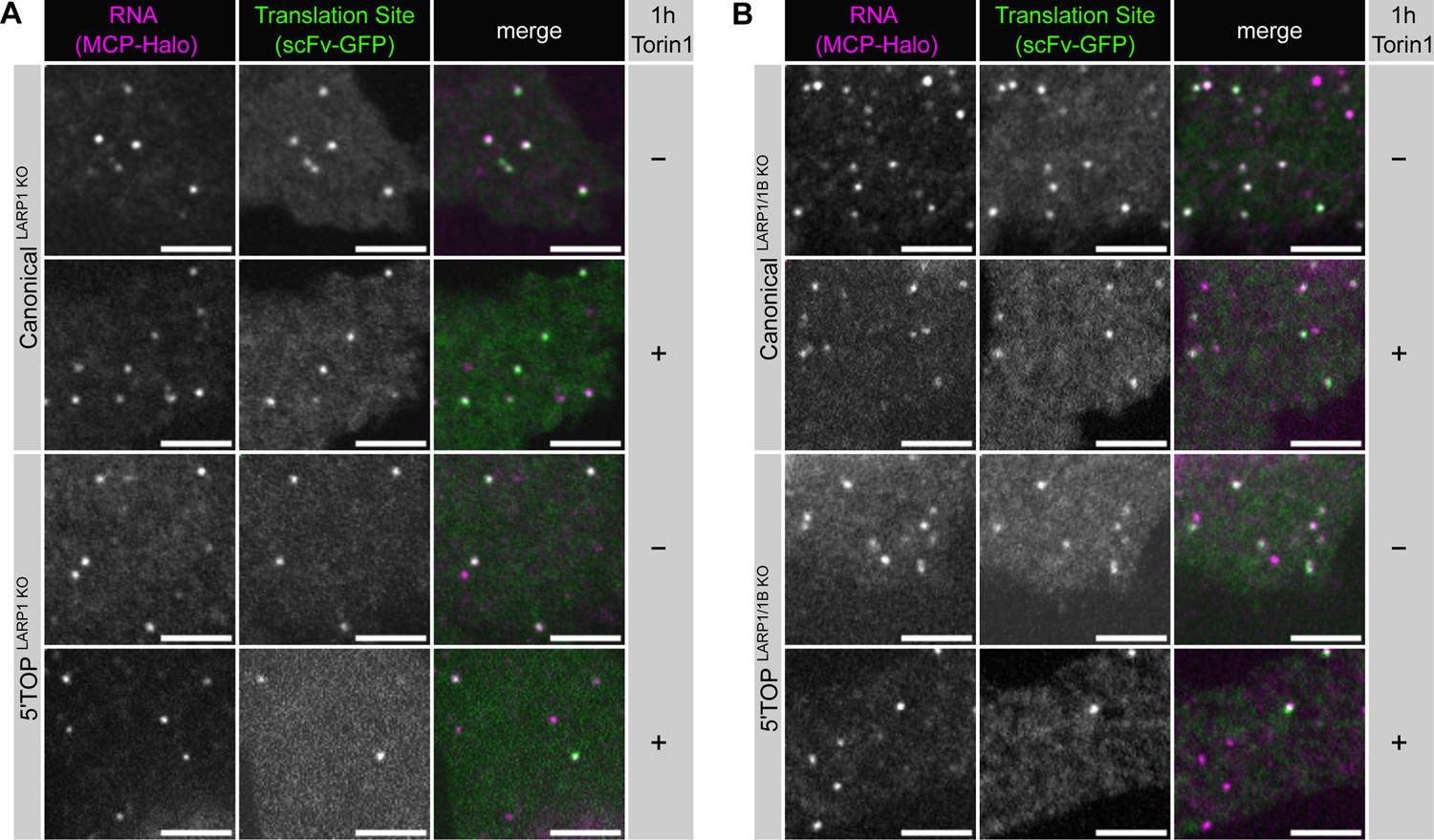
Single-molecule imaging of translation in LARP1 KO and LARP1/1B dKO cell lines. (**A**) Representative images of LARP1 KO cell lines expressing canonical and 5′TOP mRNAs (MCP-Halo foci, magenta) undergoing translation (scFv-GFP foci, green) in the absence or presence of mTOR inhibitor Torin 1 (250nM, 1h). Scale bars = 5 µm. (**B**) Representative images of LARP1/1B dKO cell lines expressing canonical and 5′TOP mRNAs (MCP-Halo foci, magenta) undergoing translation (scFv-GFP foci, green) in absence or presence of mTOR inhibitor Torin 1 (250nM, 1h). Scale bars = 5 µm.

**Fig. S7.**
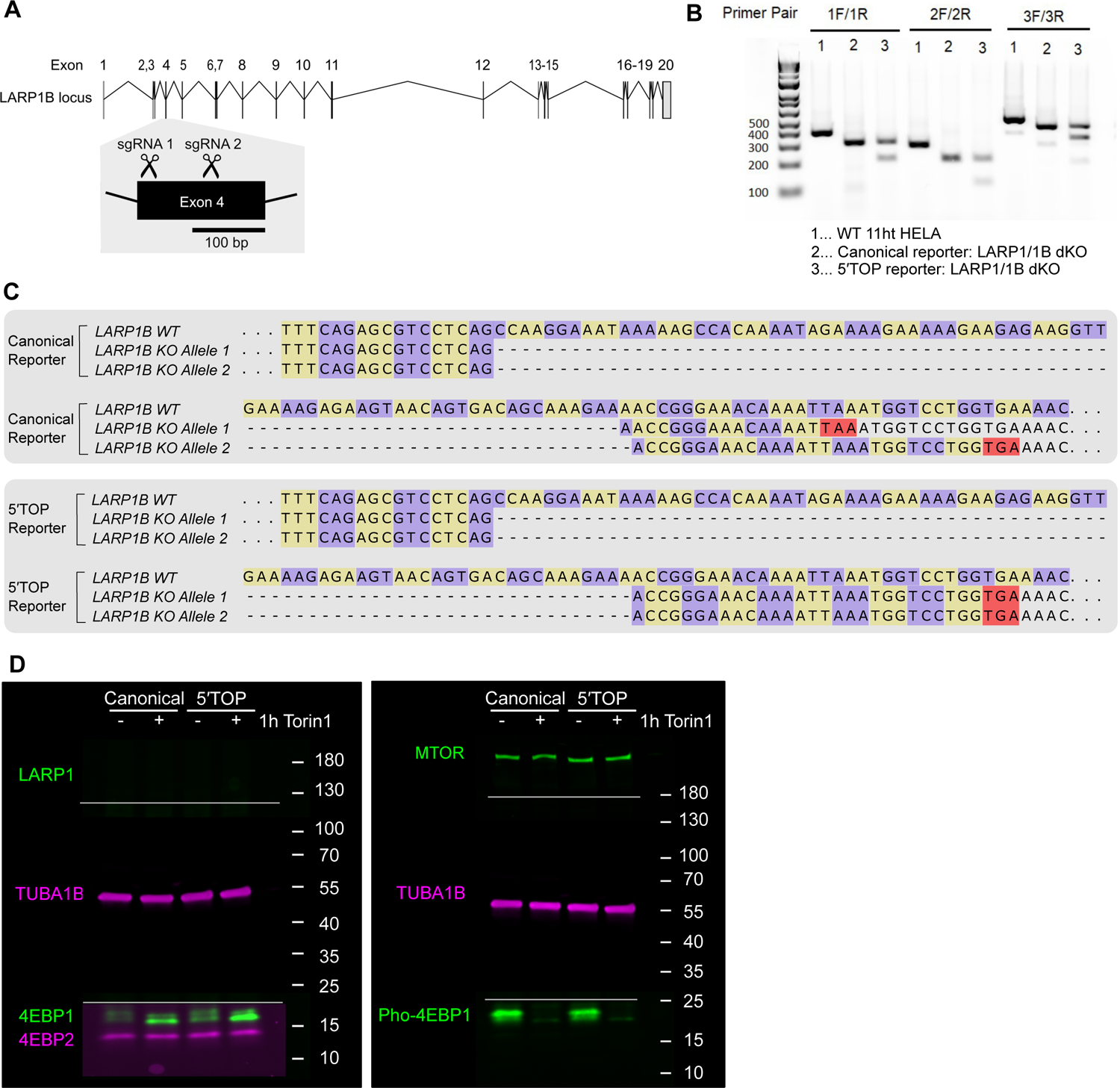
Validation of LARP1/1B dKO cell lines. (**A**) CRISPR-Cas9 editing strategy for LARP1B gene locus, using two sgRNAs targeting exon 4 of LARP1B upstream of any domain of known function (sgRNA 1 taken from (30)). (**B**) Validation of ∼100bp truncations of LARP1B, using cDNA generated from total RNA extracted from LARP1/1B dKO canonical and 5′TOP mRNA cell lines used for single-molecule imaging. (**C**) Genotyping of canonical and 5′TOP LARP1/1B dKO clones used for single-molecule imaging (Fig. 2C-E), showing ∼100 bp truncations of LARP1B exon 4 resulting in frameshifting and premature termination codons. (**D**) Western blot analysis of mTORC1 signaling in canonical and 5′TOP LARP1/1B dKO cell lines used for single-molecule imaging. Upon 1h Torin1, 4EBP1 is shifted to its dephosphorylated state, as seen by the lower migration size of 4EBP1 (4EBP1 antibody, left), and the disappearance of Phospho-4EBP1 (Phospho-Ser65-4EBP1 antibody, right). Lines indicate cut membrane pieces probed with different mouse (magenta) and rabbit (green) antibodies, and imaged together using two-color fluorescent imaging. Brightness and contrast were individually adjusted for each antibody shown.

**Fig. S8.**
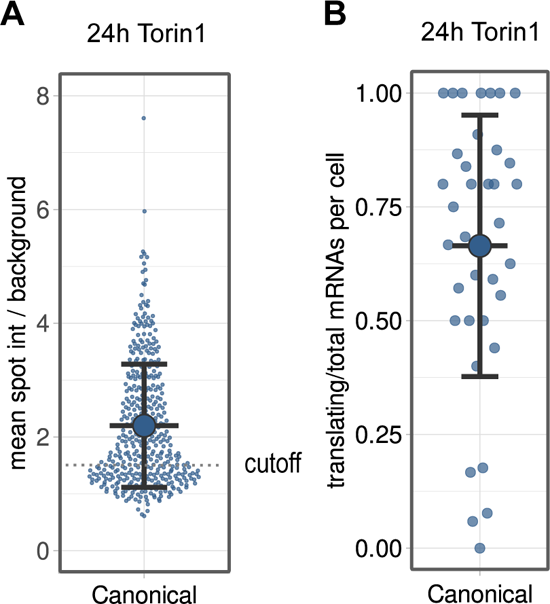
Translation of canonical mRNAs under long-term Torin1 treatment. (**A**) Quantification of SunTag signal of translation sites for canonical mRNA WT cells after 24hr Torin1 treatment (250nM). SunTag intensities are plotted for 420 mRNAs overlaid with the mean ± SD. (**B**) Fraction of mRNAs undergoing translation quantified per cell after 24 hr Torin1 treatment (250nM). Individual values are plotted overlaid with the mean ± SD of 37 cells.

**Fig. S9.**
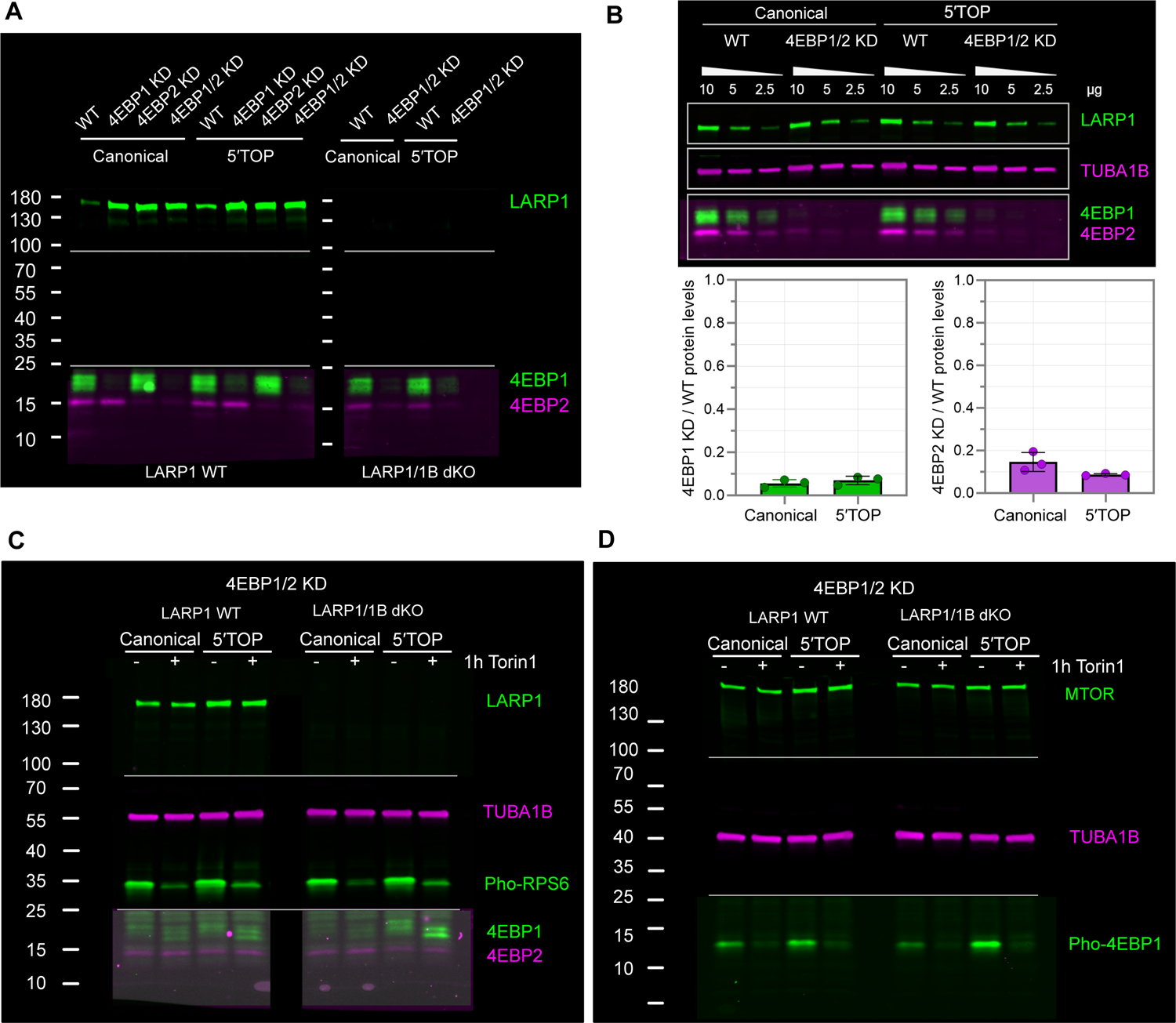
Validation of shRNA knockdown of 4EBP1/2 proteins. (**A**) Western blot analysis of shRNA-mediated stable knockdown of 4EBP1/2 proteins in canonical and 5′TOP mRNA cell lines with WT LARP1 (left) or LARP1/1B dKO (right). Depletion of 4EBP1 and 4EBP2 proteins was analyzed in mTOR active cells with 4EBP1 and 4EBP2 antibodies. (**B**) Quantification of knock-down efficiency in canonical and 5′TOP mRNA cell lines with WT LARP1. Total lysate was loaded in amounts of 10, 5, and 2.5 µg to test antibody linearity, and semi-quantitative estimates of knockdown efficiency were calculated (4EBP1 KD/WT) for both cell lines. (**C**) Western blot analysis of mTORC1 signaling in 4EBP1/2 dKD cell lines with WT LARP1 (left) or LARP1/1B dKO (right). Upon 1h Torin1, RPS6 became dephosphorylated as seen by the decreased signal for Phospho-RPS6 (Phospho-Ser235/236-RPS6 antibody). 4EBP1 and 4EBP2 are detected at low levels. (**D**) Western blot analysis of mTORC1 signaling in 4EBP1/2 dKD reporter cell lines with WT LARP1 (left) or LARP1/1B dKO (right). Upon 1h Torin1 (250nM), residual 4EBP1 becomes dephosphorylated as seen by the disappearance of Phospho-4EBP1 (Ser65). Lines indicate cut membrane pieces probed with different mouse (magenta) and rabbit (green) antibodies, and imaged together using two-color fluorescent imaging. Brightness and contrast were individually adjusted for each antibody shown.

**Fig. S10.**
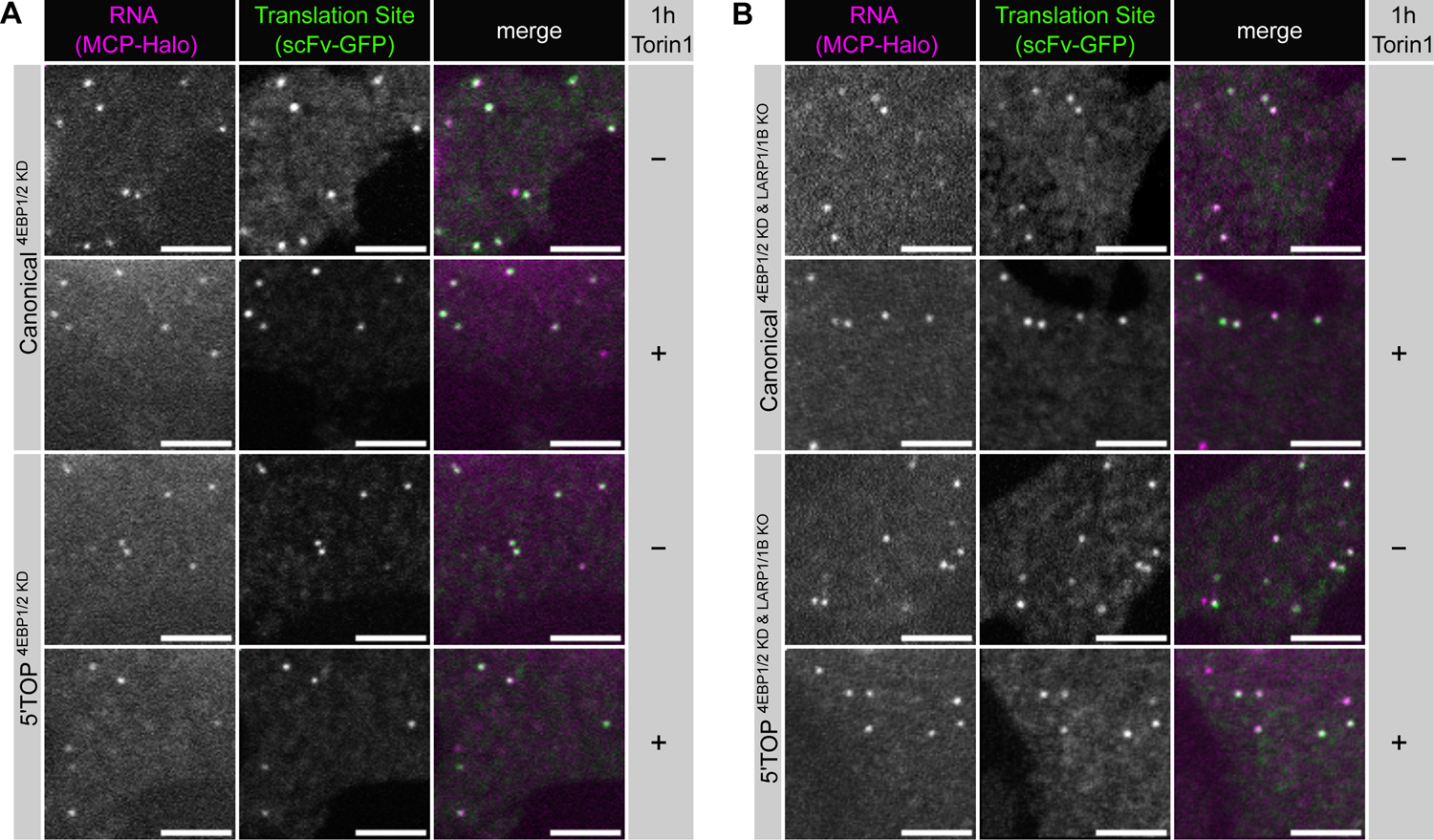
Single-molecule imaging of translation in 4EBP1/2 dKD, 4EBP1/2_ LARP1/1B dKO cell lines. (**A**) Representative images of canonical and 5′TOP reporter mRNAs in 4EBP1/2 dKD cells (MCP-Halo foci, magenta) undergoing translation (scFv-GFP foci, green) in absence or presence of mTOR inhibitor Torin 1 (250nM, 1h). Scale bars = 5 µm. (**B**) Representative images of canonical and 5′TOP mRNAs in 4EBP1/2 dKD_LARP1/1B dKO cells (MCP-Halo foci, magenta) undergoing translation (scFv-GFP foci, green) in absence or presence of mTOR inhibitor Torin 1 (250nM, 1h). Scale bars = 5 µm.

**Fig. S11.**
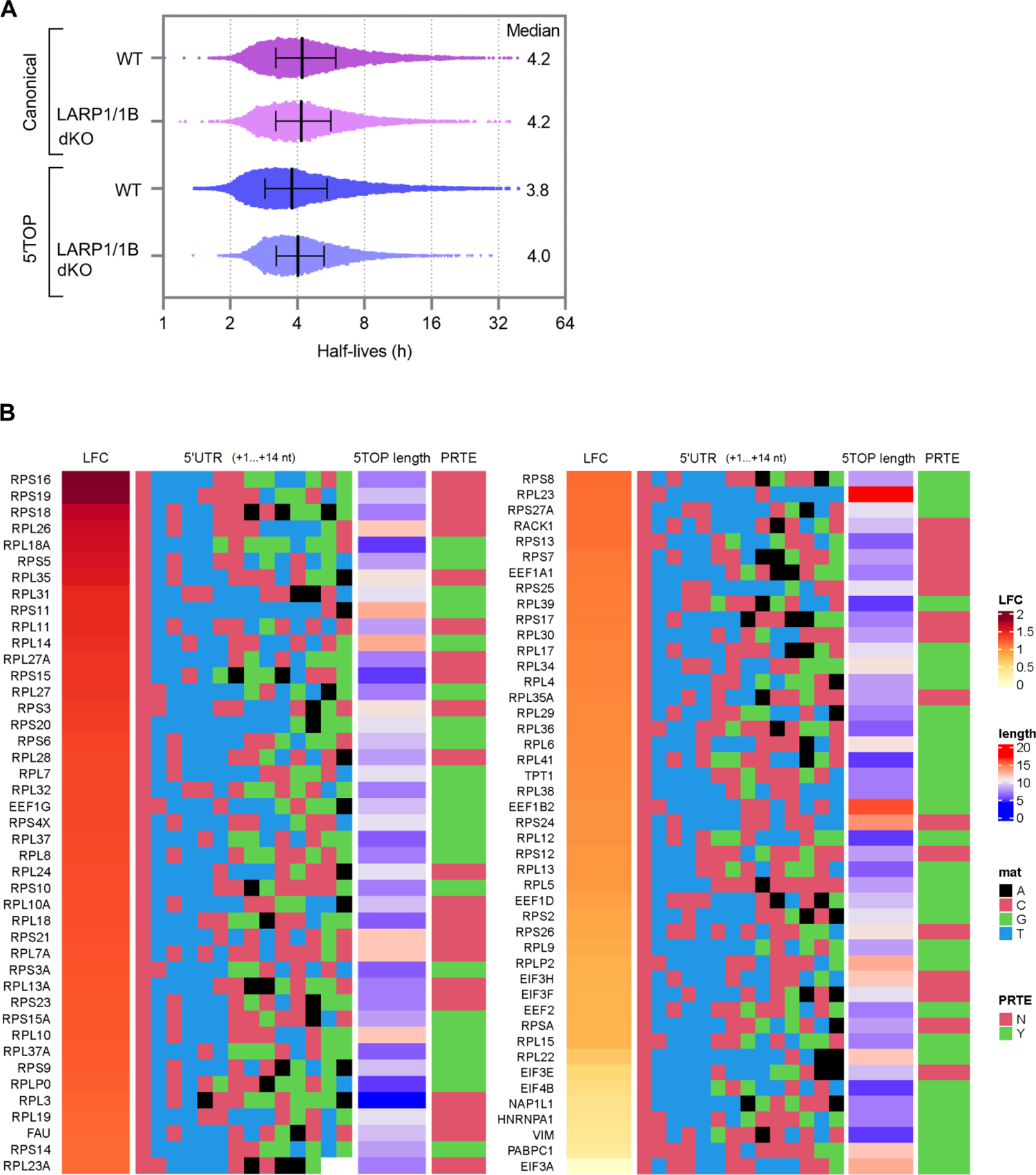
Global analysis of mRNA half-lives determined by SLAM-seq. (**A**) Distribution of RNA half-lives estimated from SLAM-seq data for WT and LARP1/1B dKO cell lines. Values are plotted with the median and interquartile range. (**B**) List of 5′TOP mRNAs from SLAMseq data, ranked by log2 fold change (LFC). The sequence of the first 14 nt of the 5′UTR containing the 5′TOP motif (RPL23A 5′UTR: 12 nt), the length of the 5′TOP motif, and presence or absence of a downstream pyrimidine-rich element (PRTE) within the 5′UTR is shown.

## Supplementary tables

**Tables S1,** S3-S5 are provided as separate files.

**Table S1. RNA-seq results of LARP1 KO cells compared to WT cells.** Differential gene expression (log2 FC) was calculated with the Bioconductor package edgeR, using an expression cut-off of log2 CPM ≥1 to exclude low confidence transcripts, and using gene type annotation to exclude pseudogenes.

**Table S2.** Reference list of 5′TOP mRNAs. Absence (0) or presence (1) of a putative pyrimidinerich translational element is listed.

| gene name | PRTE | gene name | PRTE | gene name | PRTE | gene name | PRTE |
| --- | --- | --- | --- | --- | --- | --- | --- |
| EEF1A1 | 0 | RPL13A | 0 | RPL36A | 1 | RPS18 | 0 |
| EEF1B2 | 1 | RPL14 | 1 | RPL37 | 1 | RPS19 | 0 |
| EEF1D | 1 | RPL15 | 1 | RPL37A | 1 | RPS2 | 1 |
| EEF1G | 1 | RPL17 | 1 | RPL38 | 1 | RPS20 | 1 |
| EEF2 | 1 | RPL18 | 0 | RPL39 | 1 | RPS21 | 0 |
| EIF2A | 0 | RPL18A | 1 | RPL3L | 0 | RPS23 | 0 |
| EIF2S3 | 0 | RPL19 | 0 | RPL4 | 1 | RPS24 | 0 |
| EIF3A | 1 | RPL21 | 0 | RPL41 | 1 | RPS25 | 0 |
| EIF3E | 0 | RPL22 | 1 | RPL5 | 1 | RPS26 | 0 |
| EIF3F | 0 | RPL22L1 | 0 | RPL6 | 1 | RPS27 | 0 |
| EIF3H | 0 | RPL23 | 1 | RPL7 | 1 | RPS27A | 1 |
| EIF3K | 0 | RPL23A | 0 | RPL7A | 0 | RPS28 | 0 |
| EIF3L | 0 | RPL24 | 0 | RPL8 | 1 | RPS29 | 0 |
| EIF4A2 | 0 | RPL26 | 0 | RPL9 | 1 | RPS3 | 0 |
| EIF4B | 1 | RPL27 | 1 | RPLP0 | 1 | RPS3A | 1 |
| FAU | 0 | RPL27A | 0 | RPLP1 | 0 | RPS4X | 1 |
| GNB2L1 | 1 | RPL28 | 0 | RPLP2 | 1 | RPS4Y1 | 0 |
| HNRNPA1 | 1 | RPL29 | 1 | RPS10 | 1 | RPS4Y2 | 0 |
| NAP1L1 | 1 | RPL3 | 0 | RPS11 | 1 | RPS5 | 1 |
| PABPC1 | 1 | RPL30 | 0 | RPS12 | 0 | RPS6 | 1 |
| RACK1 | 0 | RPL31 | 1 | RPS13 | 0 | RPS7 | 0 |
| RPL10 | 1 | RPL32 | 1 | RPS14 | 1 | RPS8 | 1 |
| RPL10A | 0 | RPL34 | 1 | RPS15 | 0 | RPS9 | 1 |
| RPL11 | 0 | RPL35 | 0 | RPS15A | 1 | RPSA | 0 |
| RPL12 | 1 | RPL35A | 0 | RPS16 | 0 | TPT1 | 1 |
| RPL13 | 1 | RPL36 | 1 | RPS17 | 0 | VIM | 1 |

**Table S3. RNA-seq results of 4EBP1/2 dKD cells compared to WT cells.** An expression cutoff of log2 CPM ≥1 was used to exclude low confidence transcripts, and gene type annotation to exclude pseudogenes.

**Table S4. SLAM-seq results of LARP1/1B dKO cells compared to WT cells.** Low confidence half-life estimates were excluded using a Rsquared threshold of >0.75 in all conditions, and pseudogenes were excluded using gene type annotation.

**Table S5. Reagent list.**

## References

1. Liu GY, Sabatini DM. mTOR at the nexus of nutrition, growth, ageing and disease. Nature Reviews Molecular Cell Biology. 2020;21(4):183–203.

2. Battaglioni S, Benjamin D, Wälchli M, Maier T, Hall MN. mTOR substrate phosphorylation in growth control. Cell. 2022;185(11):1814–36.

3. Jia J-J, Lahr RM, Solgaard MT, Moraes BJ, Pointet R, Yang A-D, et al. mTORC1 promotes TOP mRNA translation through site-specific phosphorylation of LARP1. Nucleic Acids Research. 2021.

4. Fonseca BD, Zakaria C, Jia JJ, Graber TE, Svitkin Y, Tahmasebi S, et al. La-related Protein 1 (LARP1) Represses Terminal Oligopyrimidine (TOP) mRNA Translation Downstream of mTOR Complex 1 (mTORC1). J Biol Chem. 2015;290(26):15996–6020.

5. Philippe L, Vasseur JJ, Debart F, Thoreen CC. La-related protein 1 (LARP1) repression of TOP mRNA translation is mediated through its cap-binding domain and controlled by an adjacent regulatory region. Nucleic Acids Res. 2018;46(3):1457–69.

6. Meyuhas O, Kahan T. The race to decipher the top secrets of TOP mRNAs. Biochimica et Biophysica Acta (BBA) - Gene Regulatory Mechanisms. 2015;1849(7):801–11.

7. Biberman Y, Meyuhas O. Substitution of just five nucleotides at and around the transcription start site of rat β-actin promoter is sufficient to render the resulting transcript a subject for translational control. FEBS Letters. 1997;405(3):333–6.

8. Avni D, Shama S, Loreni F, Meyuhas O. Vertebrate mRNAs with a 5’-terminal pyrimidine tract are candidates for translational repression in quiescent cells: characterization of the translational cis-regulatory element. Molecular and Cellular Biology. 1994;14(6):3822–33.

9. Berman AJ, Thoreen CC, Dedeic Z, Chettle J, Roux PP, Blagden SP. Controversies around the function of LARP1. RNA Biology. 2020:1–11.

10. Thoreen CC, Chantranupong L, Keys HR, Wang T, Gray NS, Sabatini DM. A unifying model for mTORC1-mediated regulation of mRNA translation. Nature. 2012;485(7396):109-13.

11. Miloslavski R, Cohen E, Avraham A, Iluz Y, Hayouka Z, Kasir J, et al. Oxygen sufficiency controls TOP mRNA translation via the TSC-Rheb-mTOR pathway in a 4E-BP-independent manner. Journal of Molecular Cell Biology. 2014;6(3):255–66.

12. Lahr RM, Fonseca BD, Ciotti GE, Al-Ashtal HA, Jia JJ, Niklaus MR, et al. La-related protein 1 (LARP1) binds the mRNA cap, blocking eIF4F assembly on TOP mRNAs. Elife. 2017;6.

13. Lindqvist L, Imataka H, Pelletier J. Cap-dependent eukaryotic initiation factor-mRNA interactions probed by cross-linking. Rna. 2008;14(5):960–9.

14. Tamarkin-Ben-Harush A, Vasseur JJ, Debart F, Ulitsky I, Dikstein R. Cap-proximal nucleotides via differential eIF4E binding and alternative promoter usage mediate translational response to energy stress. Elife. 2017;6.

15. Schofield JA, Duffy EE, Kiefer L, Sullivan MC, Simon MD. TimeLapse-seq: adding a temporal dimension to RNA sequencing through nucleoside recoding. Nature Methods. 2018;15(3):221–5.

16. Herzog VA, Reichholf B, Neumann T, Rescheneder P, Bhat P, Burkard TR, et al. Thiol-linked alkylation of RNA to assess expression dynamics. Nature Methods. 2017;14(12):1198–204.

17. Kozlov G, Mattijssen S, Jiang J, Nyandwi S, Sprules T, Iben James R, et al. Structural basis of 3′-end poly(A) RNA recognition by LARP1. Nucleic Acids Research. 2022;50(16):9534–47.

18. Mattijssen S, Kozlov G, Gaidamakov S, Ranjan A, Fonseca BD, Gehring K, et al. The isolated La-module of LARP1 mediates 3’ poly(A) protection and mRNA stabilization, dependent on its intrinsic PAM2 binding to PABPC1. RNA Biology. 2021;18(2):275–89.

19. Ogami K, Oishi Y, Sakamoto K, Okumura M, Yamagishi R, Inoue T, et al. mTOR- and LARP1-dependent regulation of TOP mRNA poly(A) tail and ribosome loading. Cell reports. 2022;41(4):111548.

20. Ogami K, Oishi Y, Nogimori T, Sakamoto K, Hoshino S-i. LARP1 facilitates translational recovery after amino acid refeeding by preserving long poly(A)-tailed TOP mRNAs. bioRxiv. 2019:716217.

21. Smith EM, Benbahouche Nour El H, Morris K, Wilczynska A, Gillen S, Schmidt T, et al. The mTOR regulated RNA-binding protein LARP1 requires PABPC1 for guided mRNA interaction. Nucleic Acids Research. 2020;49(1):458-78.

22. Hong S, Freeberg MA, Han T, Kamath A, Yao Y, Fukuda T, et al. LARP1 functions as a molecular switch for mTORC1-mediated translation of an essential class of mRNAs. Elife. 2017;6.

23. Mura M, Hopkins TG, Michael T, Abd-Latip N, Weir J, Aboagye E, et al. LARP1 post-transcriptionally regulates mTOR and contributes to cancer progression. Oncogene. 2015;34(39):5025–36.

24. Park J, Kim M, Yi H, Baeg K, Choi Y, Lee Y-s, et al. Short poly(A) tails are protected from deadenylation by the LARP1–PABP complex. Nature Structural & Molecular Biology. 2023;30(3):330–8.

25. Wilbertz JH, Voigt F, Horvathova I, Roth G, Zhan Y, Chao JA. Single-Molecule Imaging of mRNA Localization and Regulation during the Integrated Stress Response. Molecular Cell. 2019;73(5):946–58.e7.

26. Tanenbaum ME, Gilbert LA, Qi LS, Weissman JS, Vale RD. A protein-tagging system for signal amplification in gene expression and fluorescence imaging. Cell. 2014;159(3):635–46.

27. Banaszynski LA, Chen L-c, Maynard-Smith LA, Ooi AGL, Wandless TJ. A Rapid, Reversible, and Tunable Method to Regulate Protein Function in Living Cells Using Synthetic Small Molecules. Cell. 2006;126(5):995–1004.

28. Carninci P, Sandelin A, Lenhard B, Katayama S, Shimokawa K, Ponjavic J, et al. Genome-wide analysis of mammalian promoter architecture and evolution. Nat Genet. 2006;38(6):626–35.

29. Wilbertz JH, Voigt F, Horvathova I, Roth G, Zhan Y, Chao JA. Single-Molecule Imaging of mRNA Localization and Regulation during the Integrated Stress Response. Mol Cell. 2019;73(5):946–58.e7.

30. Philippe L, van den Elzen AMG, Watson MJ, Thoreen CC. Global analysis of LARP1 translation targets reveals tunable and dynamic features of 5′ TOP motifs. Proceedings of the National Academy of Sciences. 2020;117(10):5319–28.

31. Aoki K, Adachi S, Homoto M, Kusano H, Koike K, Natsume T. LARP1 specifically recognizes the 3′ terminus of poly(A) mRNA. FEBS Letters. 2013;587(14):2173–8.

32. Al-Ashtal HA, Rubottom CM, Leeper TC, Berman AJ. The LARP1 La-Module recognizes both ends of TOP mRNAs. RNA Biology. 2019:1–11.

33. Truitt ML, Conn CS, Shi Z, Pang X, Tokuyasu T, Coady AM, et al. Differential Requirements for eIF4E Dose in Normal Development and Cancer. Cell. 2015;162(1):59–71.

34. Thoreen CC. The molecular basis of mTORC1-regulated translation. Biochemical Society Transactions. 2017;45(1):213–21.

35. Gandin V, Masvidal L, Hulea L, Gravel SP, Cargnello M, McLaughlan S, et al. nanoCAGE reveals 5’ UTR features that define specific modes of translation of functionally related MTOR-sensitive mRNAs. Genome Res. 2016;26(5):636–48.

36. van den Elzen AMG, Watson MJ, Thoreen CC. mRNA 5’ terminal sequences drive 200-fold differences in expression through effects on synthesis, translation and decay. PLoS Genet. 2022;18(11):e1010532.

37. Meyuhas O. Synthesis of the translational apparatus is regulated at the translational level. Eur J Biochem. 2000;267(21):6321–30.

38. Mamane Y, Petroulakis E, Martineau Y, Sato TA, Larsson O, Rajasekhar VK, et al. Epigenetic activation of a subset of mRNAs by eIF4E explains its effects on cell proliferation. PLoS One. 2007;2(2):e242.

39. Kajjo S, Sharma S, Brothers RW, Delisle V, Fabian RM. PABPC plays a critical role in establishing mTORC-mediated translational control. Under Preparation. 2023.

40. Tomoo K, Shen X, Okabe K, Nozoe Y, Fukuhara S, Morino S, et al. Crystal structures of 7-methylguanosine 5’-triphosphate (m(7)GTP)- and P(1)-7-methylguanosine-P(3)-adenosine-5’,5’-triphosphate (m(7)GpppA)-bound human full-length eukaryotic initiation factor 4E: biological importance of the C-terminal flexible region. Biochem J. 2002;362(Pt 3):539–44.

41. Furic L, Rong L, Larsson O, Koumakpayi IH, Yoshida K, Brueschke A, et al. eIF4E phosphorylation promotes tumorigenesis and is associated with prostate cancer progression. Proc Natl Acad Sci U S A. 2010;107(32):14134–9.

42. Zinshteyn B, Rojas-Duran MF, Gilbert WV. Translation initiation factor eIF4G1 preferentially binds yeast transcript leaders containing conserved oligo-uridine motifs. Rna. 2017;23(9):1365–75.

43. Jin H, Xu W, Rahman R, Na D, Fieldsend A, Song W, et al. TRIBE editing reveals specific mRNA targets of eIF4E-BP in <em>Drosophila</em> and in mammals. Science Advances. 2020;6(33):eabb8771.

44. Fuentes P, Pelletier J, Martinez-Herráez C, Diez-Obrero V, Iannizzotto F, Rubio T, et al. The 40S-LARP1 complex reprograms the cellular translatome upon mTOR inhibition to preserve the protein synthetic capacity. Sci Adv. 2021;7(48):eabg9275.

45. Gentilella A, Moron-Duran FD, Fuentes P, Zweig-Rocha G, Riano-Canalias F, Pelletier J, et al. Autogenous Control of 5’TOP mRNA Stability by 40S Ribosomes. Mol Cell. 2017;67(1):55–70 e4.

46. Brito Querido J, Sokabe M, Kraatz S, Gordiyenko Y, Skehel JM, Fraser CS, et al. Structure of a human 48S translational initiation complex. Science. 2020;369(6508):1220-7.

47. Goering R, Arora A, Pockalny MC, Taliaferro JM. RNA localization mechanisms transcend cell morphology. eLife. 2023;12:e80040.

48. Livingston NM, Kwon J, Valera O, Saba JA, Sinha NK, Reddy P, et al. Bursting Translation on Single mRNAs in Live Cells. bioRxiv. 2022:2022.11.07.515520.

49. Dave P, Roth G, Griesbach E, Mateju D, Hochstoeger T, Chao JA. Single-molecule imaging reveals translation-dependent destabilization of mRNAs. Molecular Cell. 2023;83(4):589–606.e6.

50. Cong L, Ran FA, Cox D, Lin S, Barretto R, Habib N, et al. Multiplex genome engineering using CRISPR/Cas systems. Science. 2013;339(6121):819-23.

51. Chu VT, Weber T, Wefers B, Wurst W, Sander S, Rajewsky K, et al. Increasing the efficiency of homology-directed repair for CRISPR-Cas9-induced precise gene editing in mammalian cells. Nat Biotechnol. 2015;33(5):543–8.

52. Bryant DM, Datta A, Rodríguez-Fraticelli AE, Peränen J, Martín-Belmonte F, Mostov KE. A molecular network for de novo generation of the apical surface and lumen. Nat Cell Biol. 2010;12(11):1035–45.

53. Schindelin J, Arganda-Carreras I, Frise E, Kaynig V, Longair M, Pietzsch T, et al. Fiji: an open-source platform for biological-image analysis. Nature methods. 2012;9(7):676–82.

54. Rueden CT, Schindelin J, Hiner MC, DeZonia BE, Walter AE, Arena ET, et al. ImageJ2: ImageJ for the next generation of scientific image data. BMC bioinformatics. 2017;18(1):1–26.

55. Preibisch S, Saalfeld S, Schindelin J, Tomancak P. Software for bead-based registration of selective plane illumination microscopy data. Nature methods. 2010;7(6):418–9.

56. Mateju D, Eichenberger B, Voigt F, Eglinger J, Roth G, Chao JA. Single-Molecule Imaging Reveals Translation of mRNAs Localized to Stress Granules. Cell. 2020;183(7):1801–12.e13.

57. Tinevez J-Y, Perry N, Schindelin J, Hoopes GM, Reynolds GD, Laplantine E, et al. TrackMate: An open and extensible platform for single-particle tracking. Methods. 2017;115:80–90.

58. Lord SJ, Velle KB, Mullins RD, Fritz-Laylin LK. SuperPlots: Communicating reproducibility and variability in cell biology. Journal of Cell Biology. 2020;219(6).

59. Hsieh AC, Liu Y, Edlind MP, Ingolia NT, Janes MR, Sher A, et al. The translational landscape of mTOR signalling steers cancer initiation and metastasis. Nature. 2012;485(7396):55-61.

60. Gaidatzis D, Lerch A, Hahne F, Stadler MB. QuasR: quantification and annotation of short reads in R. Bioinformatics. 2014;31(7):1130–2.

61. Neumann T, Herzog VA, Muhar M, von Haeseler A, Zuber J, Ameres SL, et al. Quantification of experimentally induced nucleotide conversions in high-throughput sequencing datasets. BMC Bioinformatics. 2019;20(1):258.

62. Danecek P, Auton A, Abecasis G, Albers CA, Banks E, DePristo MA, et al. The variant call format and VCFtools. Bioinformatics. 2011;27(15):2156–8.

63. Lun AT, Chen Y, Smyth GK. It’s DE-licious: A Recipe for Differential Expression Analyses of RNA-seq Experiments Using Quasi-Likelihood Methods in edgeR. Methods Mol Biol. 2016;1418:391–416.

